# Goofy/123Cre Lineage Tracing Differentiates Olfactory and Vomeronasal Neurons from GnRH-1 and Terminal Nerve Neurons During Neuronal Migration and Reveals Additional Olfactory Placode-Derived Cells in the Brain

**DOI:** 10.64898/2025.12.17.694945

**Authors:** Enrico Amato, Mia V. Call, Noah M. LeFever, Mya Aviles-Carlos, Nikki M. Dolphin, Paolo E. Forni

## Abstract

The olfactory placode (OP) gives rise to a wide array of chemosensory neurons in the nasal region, including olfactory sensory neurons, vomeronasal sensory neurons, and those of the septal organ and Grueneberg ganglion. During placodal invagination, the OP also produces migratory neurons, including gonadotropin-releasing hormone-1 (GnRH-1) neurons, somatostatin-expressing neurons, Prokineticin Receptor 2 (Prokr2) pioneer/terminal nerve (TN) neurons, which are thought to initiate olfactory bulb development. Despite decades of research, the genetic lineage and molecular identity of OP-derived neuronal types remain under investigation. GnRH-1 neurons play essential roles in reproduction and chemodetection but appear genetically distinct from olfactory and vomeronasal chemosensory neurons. The Golgi-associated olfactory signaling regulator Goofy/Gfy is broadly expressed across the placode-derived nasal chemosensory neurons. To determine whether its expression is specific to nasal chemosensory neurons or it is a shared genetic feature across OP derivatives we characterized Gfy expression and lineage at embryonic and postnatal stages. Our results confirm broad Gfy expression in developing chemosensory neurons and subsets of olfactory pioneer/TN neurons. However, we found that while Gfy was not expressed in GnRH-1 neurons while migrating, analysis at late development and postnatal stages, revealed the existence of Gfy-traced neuronal populations in the basal forebrain, some of which also express GnRH. These findings uncover previously unrecognized genetic heterogeneity among migratory nasal neurons and reinforce previous studies suggesting the existence of additional neurons with nasal origin in the brain. In addition to this, analysis of Gfy expression along the developmental trajectory of vomeronasal sensory neurons at postnatal stages revealed intriguing differences in the developmental dynamics across the two main types of vomeronasal sensory neurons.

## Introduction

The olfactory placode (OP) is an ectodermal thickening that gives rise to the vertebrate olfactory system (Verwoerd and van Oostrom, 1979). In mice, the OP primordium is first identifiable around embryonic day (E) 9.0 (Ikeda et al., 2007; Klein and Graziadei, 1983; Rawson, 1999). By E10.5, the OP invaginates to form the olfactory pit (Brunjes and Frazier, 1986; Tarozzo et al., 1995), which subsequently generates multiple specialized chemosensory structures, including the main olfactory epithelium (OE), vomeronasal organ (VNO), septal organ, and Gruenberg ganglion (GG) (Couly and Le Douarin, 1985; Cuschieri, 1975).

As the OP forms and invaginates, it undergoes successive waves of neurogenesis (Forni et al., 2013; Paronett et al., 2023; Rukh et al., 2024). Shortly after its formation, between E10.5 and E11.5, the OP is highly proliferative and displays abundant mitotic figures within its apical layers (Corepal et al., 2019; Cuschieri, 1975; Graham et al., 2007; Taroc et al., 2020a). During this early window, the OP primarily generates migratory neurons that delaminate and move toward the developing forebrain (Fornaro et al., 2003; Forni et al., 2011; Kawauchi et al., 2004; Lassiter et al., 2014; Murakami et al., 2022). These early migratory populations include gonadotropin-releasing hormone-1 (GnRH-1) neurons, presumptive terminal nerve (TN) neurons, and olfactory pioneer neurons, which are essential for olfactory bulb (OB) induction (Chen et al., 2026; Gong and Shipley, 1995).

Around E11, neurogenesis within the olfactory epithelia shifts toward the production of chemosensory neurons. This transition coincides with a marked increase in Achaete-scute homolog-1 (Ascl1)-expressing neuronal progenitors and the establishment of neurogenesis for the sensory neurons of the OE and VNO (Guillemot et al., 1993; Taroc et al., 2020a). Thus, migratory neurons and chemosensory neurons emerge from temporally and molecularly distinct neurogenic programs within the OP.

GnRH-1 neurons are widely accepted to be placode-derived in vertebrates (Murakami et al., 2022; Schwanzel-Fukuda and Pfaff, 1989; Whitlock and Westerfield, 2000; Wray et al., 1989). These neurons represent an evolutionarily ancient cell population (Herbison, 2025), as GnRH peptides and GnRH-expressing neurons have been identified in ascidians, organisms that lack specialized olfactory epithelia and instead possess proto-placodal structures (Abitua et al., 2015; Eura et al., 2020; Kamiya et al., 2014; Poncelet and Shimeld, 2020; Stolfi et al., 2015). In vertebrates, GnRH-1 neurons migrate as part of a larger migratory mass that also includes TN neurons, olfactory pioneer neurons, neuropeptide Y (Npy)-expressing neurons, somatostatin (Sst)-expressing neurons, and olfactory ensheathing cells (Demski and Schwanzel-Fukuda, 1987; Hilal et al., 1996; Miller et al., 2010; Murakami et al., 2022; Taroc et al., 2017; Valverde et al., 1993; Vilensky, 2012).

GnRH-1 neurons, TN neurons, and olfactory pioneer neurons represent some of the earliest neuronal populations to differentiate in the developing nose of rodents. Prokr2Cre-based lineage tracing revealed that pioneer/TN cells can be identified between E11.5 and E14.5 by strong immunoreactivity for Contactin-2 (Cntn2/TAG-1) and Map2, markers largely absent from developing olfactory and vomeronasal sensory neurons (Amato et al., 2025b; Chen et al., 2026; Schwarting et al., 2001; Taroc et al., 2017).

Despite their common placodal origin, the nasal migratory neurons differ molecularly and genetically from the chemosensory neurons of the OE and VNO. GnRH-1 neurons uniquely express the transcription factor Isl1, defining a distinct genetic lineage within the migratory population (Lund et al., 2020; Shan et al., 2020; Taroc et al., 2020a). Consistently, GnRH-1 neurogenesis is preserved in Gli3 loss-of-function mutants, whereas olfactory and vomeronasal neurogenesis is severely disrupted (Taroc et al., 2020b). Further lineage-tracing studies using Peripherin BAC transgenic mice and Prokineticin receptor 2-Cre (Prokr2Cre) lines support the existence of genetically distinct lineages between GnRH-1 neurons and other migratory neurons, including pioneer and TN neurons (Amato et al., 2024; Taroc et al., 2017).

The functional relevance of early migratory neurons in the nose is underscored by human pathology. Patients with Kallmann syndrome exhibit anosmia or hyposmia accompanied by delayed or absent puberty (Forni and Wray, 2015; Kallmann FJ, 1944; Lieblich et al., 1982).

Beyond their canonical role in regulating the hypothalamic–pituitary–gonadal axis, GnRH-expressing neurons have been proposed to contribute directly to chemosensory processing from ascidians to vertebrates (Abitua et al., 2015; Umatani and Oka, 2019). Recent work has identified a subpopulation of GnRH-1 neurons near the OB that participates in the processing of chemosensory signals relevant to mating behavior and opposite-sex odor preference (Decoster et al., 2024; Umatani and Oka, 2019).

Despite decades of investigation, the developmental relationship between migratory neurons and the specialized chemosensory neurons of the olfactory system remains fuzzy. Progress has been limited by the scarcity of early markers that allow for the discrimination of these cell types. The Golgi-associated olfactory signaling regulator Goofy (Gfy) is an integral Golgi membrane protein expressed in nasal chemosensory neurons, including those of the OE, vomeronasal sensory epithelium (VSE), and GG (Bao et al., 2025; Enomoto et al., 2019; Kaneko-Goto et al., 2013; Lu and Matsunami, 2025). Here, we characterize Gfy expression and genetic lineage from E11.5 to postnatal stages. Moreover, we characterized Gfy expression across the main types of vomeronasal sensory neurons at postnatal stages. Our findings reveal molecular and developmental distinctions between the GnRH neurons, TN neurons, and neurons belonging to the specialized chemosensory system of rodents.

## Results

### Gfy is heterogeneously expressed across newly formed neurons in the developing olfactory system

The developing OP and its derivatives are highly neurogenic, resulting in continuous and asynchronous neuronal production throughout the nasal region. In mouse, neurogenesis in the OP starts soon after its formation around E9.5, as pioneer neurons can be already detected at E10.5 (Chen et al., 2009; Fornaro et al., 2003; Forni et al., 2011; Katoh et al., 2011). By analyzing single-cell RNA sequencing data of the nasal area at E14.5 (Fig. 1A) (Amato et al., 2024), we followed Gfy expression (Fig. 1A-I,K) and its correlation (Fig. 1J) with markers of neurogenesis and olfactory maturation. This highlighted that Gfy is not expressed by proliferative (Mki67+) olfactory-vomeronasal progenitors (Ascl1+) (Fig. 1B-C,K), nor by olfactory-vomeronasal precursors expressing Neurod1 and Neurog1 (Fig. 1D-E,K). However, its expression correlates with the RNA binding protein Elav3 (HuC), which is a pan-neuronal marker (Fig. 1F,K) and reaches its highest correlation with Gap43 (Growth Associated Protein 43), a neuron-specific protein expressed during axon growth (Karns et al., 1987; Verhaagen et al., 1989) (Fig. 1H, J).

**Figure 1.**
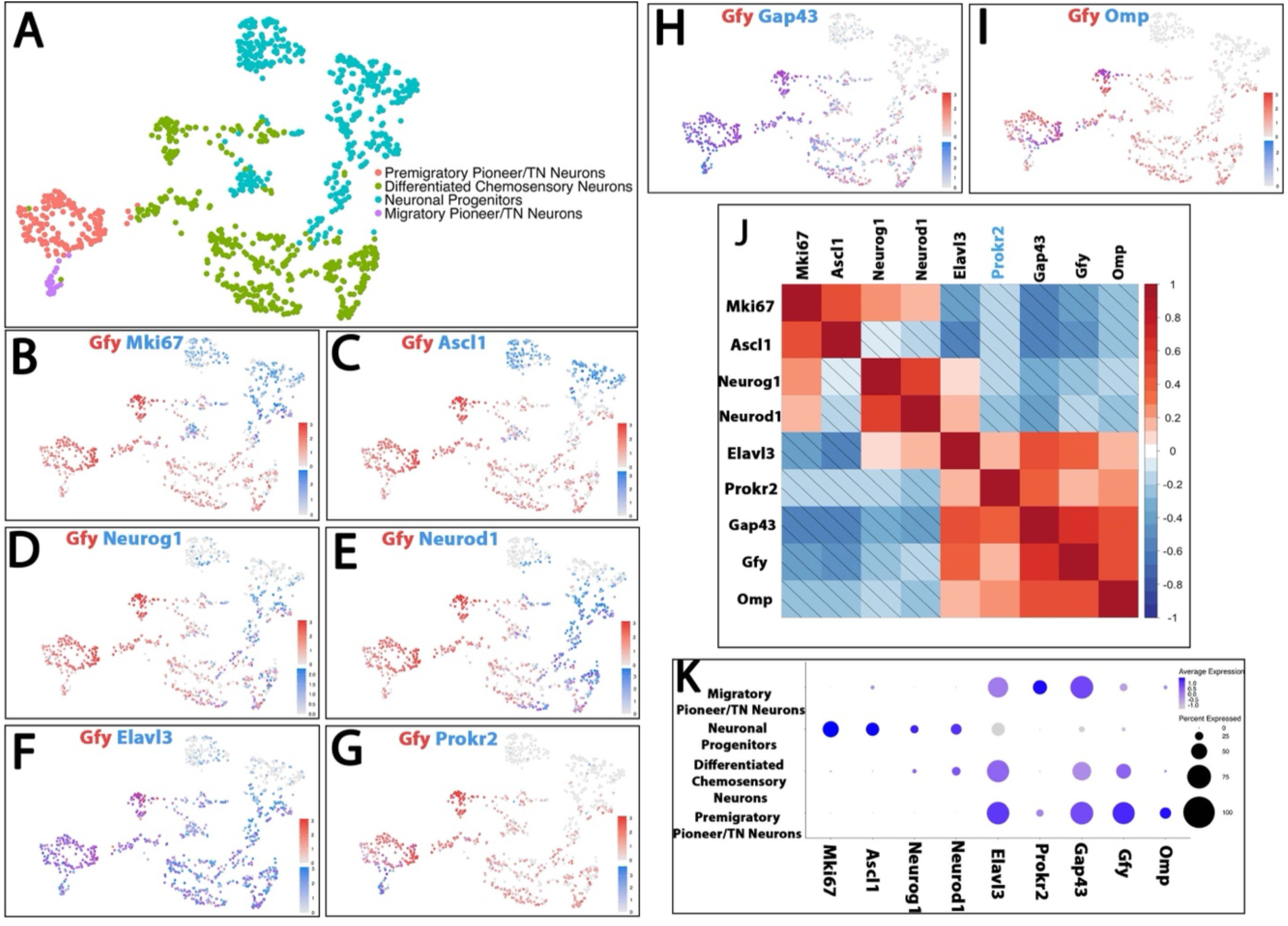
Gfy expression in the developing olfactory system. A) UMAP of neuronal populations in the developing olfactory system at E14.5 generated from single-cell RNA sequencing of the nasal area. B–I) Co-expression feature plots showing expression of Gfy with: Ascl1, Neurog1, Neurod1, Elav3, Prokr2, Gap43, and Omp expression. Progenitor markers (Ascl1, Neurog1) precede neuronal differentiation markers (Neurod1, Elavl3), while Gap43 labels immature neurons, Omp marks mature sensory neurons and Prokr2 labels pioneer/terminal nerve neurons. Gfy is enriched in differentiating and immature neuronal populations. J) Correlation matrix showing negative mRNA expression correlations (diagonal lines) between progenitor and maturation markers and positive mRNA expression correlations between Gfy and neuronal differentiation genes. K) Dotplot showing expression of progenitor and maturation markers during embryonic development of the olfactory system.

Gfy expression is retained in cell clusters containing the putative developing chemosensory neurons (OSNs/VSNs) which express the olfactory maturation marker gene Omp (Olfactory marker protein) (Fig. 1I-J). We also found lower levels of Gfy expression and correlation in the putative migratory pioneer/TN neurons (positive for Prokr2) (Amato et al., 2024) (Fig. 1G,J,K).These data support variable Gfy expression across multiple postmitotic maturing neuronal types originating from the OP (Kaneko-Goto et al., 2013; Margolis, 1972; Margolis and Tarnoff, 1973).

### The GfyCre (123Cre) mouse line can produce diCerent recombination patterns

To further characterize the Gfy genetic lineage across the various neurons originating from the OP, we bred GfyCre mice with homozygous R26tdTomato (Ai14^-/-^) reporter mice. PCR analysis of the pups showed Cre inheritance following the expected Mendelian ratios (data not shown); however (Fig. S1), among the Cre-positive pups, we observed two recombination patterns. The earliest analysis we performed is in embryos around E11 (Fig. S1A-B’). At this stage, and later stages, we found: 1) a highly chimeric recombination throughout the body, including the brain, nasal mesenchyme, olfactory epithelia, and skin; 2) a very defined recombination limited to the chemosensory regions of the developing nose. Notably, similar recombination patterns were seen regardless of whether the Cre was inherited maternally or paternally. In mice with broad recombination, the reporter could be detected before Gfy recombination in the olfactory epithelium, suggesting recombination at an early embryonic developmental stage (Fig. S1A-E). The different recombination pattern was easily identifiable by illuminating the whole embryo at the stereomicroscope before embedding (Fig. S1D). From this point on, we will discuss only results from animals in which recombination is limited to the nasal chemosensory epithelia.

We examined embryos at multiple stages from E11.5 to E18.5 (Fig. 2; Fig. 5). Although Gfy mRNA expression could be detected across developing olfactory neurons and nasal migratory neurons (Fig. 1; Fig. S2A-C’’), between E11 and E11.5, only a few neurons positive for recombination could be identified, if any (Fig. 2A). However, at E12.5 and E13.5, neurons positive for recombination were more clearly identifiable in the developing chemosensory epithelia and as part of the migratory mass associated with Peripherin-positive fibers (Fig. 2B-C). By E14.5 and E15.5, in line with previous reports, widespread recombination was evident across the olfactory-vomeronasal epithelia and their projections to the OB (Fig. 2D-E), as well as in neurons of the Gruenberg ganglion (GG) (Fig. S3) (Kaneko-Goto et al., 2013).

**Figure 2.**
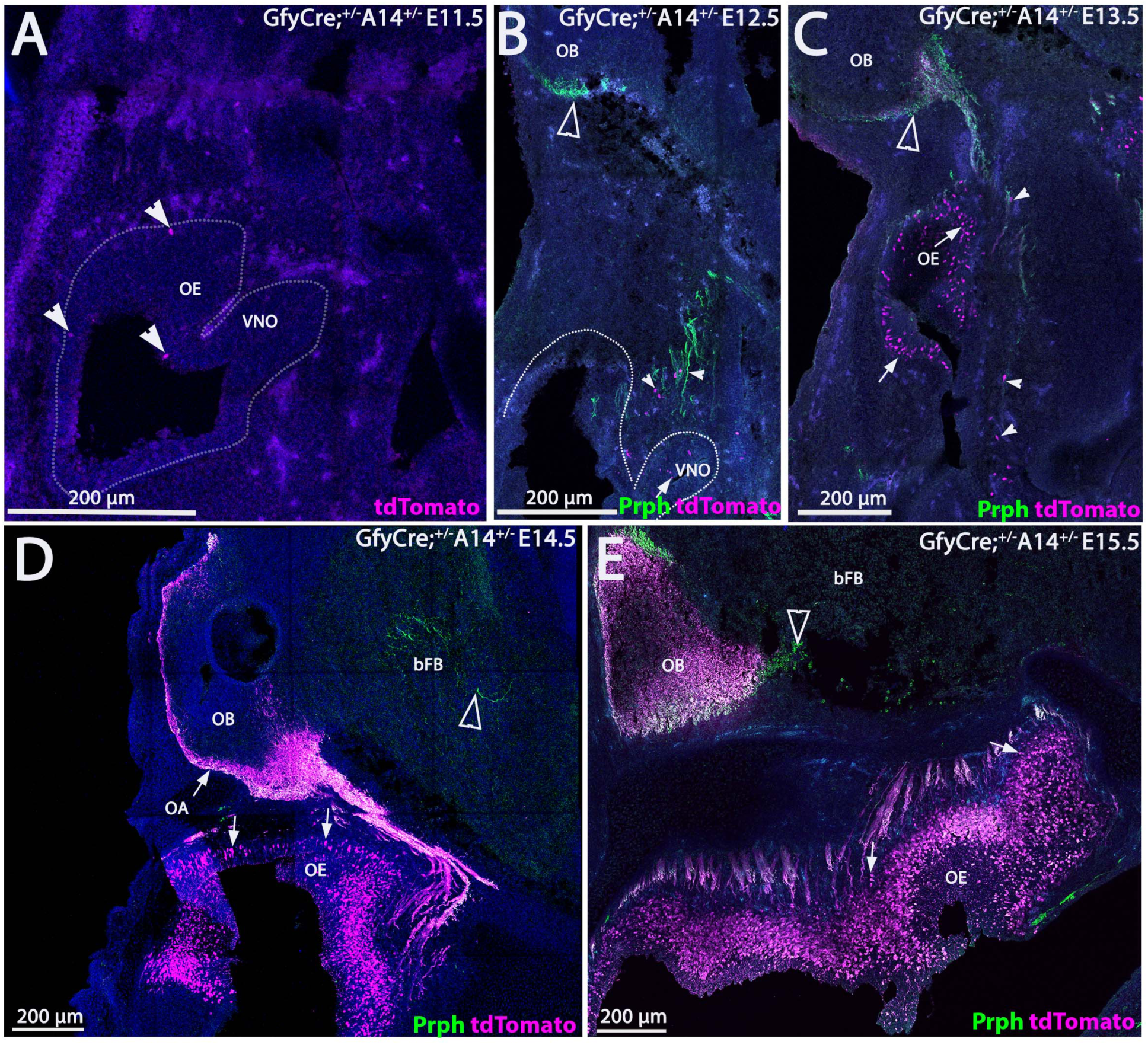
Time course of Gfy genetic lineage tracing in the developing olfactory system. A) At E11.5, GfyCre^+/−^;Ai14^+/−^ parasagittal sections show sparse Gfy lineage tracing (magenta, arrowheads) in the olfactory epithelium (OE) and vomeronasal organ (VNO). B) At E12.5, sparse Gfy tracing is detected in both chemosensory neurons (tailed arrows) and migratory neurons (white notched arrowheads), closely associated with peripherin-positive fibers (Prph, green), including near the olfactory bulb (OB). C) At E13.5, Gfy-traced cells are present in the OE (tailed arrows) and within the peripherin-positive migratory mass (notched arrowheads); peripherin-positive axons positive for the tracing are visible near the OB (open arrowhead). D–E) At E14.5 and E15.5, Gfy tracing becomes abundant throughout the nasal epithelium, in the OE and olfactory axons (OA) surrounding the OB (tailed arrows). In contrast, Peripherin-positive axons projecting to the basal forebrain (bFB) are largely Gfy-negative (open arrowheads). Scale bars A-E, 200 µm.

Notably, at E14.5 and E15.5, we also observed Peripherin-positive fibers resembling those of the putative TN (Chen et al., 2026) invading the brain ventrally to the OB, which appeared negative for recombination (Fig. 2D-E) (Fueshko and Wray, 1994; Wray et al., 1994).

### In developing mice, Gfy recombination is detected 24-48 hours after neurogenesis

The migratory neurons forming in the OP are readily detectable at E10.5 (Fornaro et al., 2003; Forni et al., 2011; Miller et al., 2010), prior to the initial neurogenesis of the olfactory/vomeronasal chemosensory neurons that begins around E11.5 (Rawson et al., 2010; Taroc et al., 2020b). Around 2-3 days after the beginning of neurogenesis in the OP (E11.5 and E12.5), the vomeronasal epithelium and part of the migratory mass showed only sparse GfyCre recombination (Fig. 2A-B).

To get a time reference between the birth of chemosensory neurons and GfyCre recombination, we first injected GfyCre^+/-^/A14^-/-^ embryos with EdU at E10.5, then again at E11.5, and analyzed them at E14.5. (Fig. 3A). At this stage, we found that only ∼60% (SD ± 3.02%) of EdU-positive neurons born 3 or 4 days before showed detectable recombination (Fig. 3B-E). The neuronal marker HuCD was used to confirm that EdU labelled cells were neurons for this quantification (Fig. S4). EdU injections at E10.5 and E11.5 revealed that 50% (SD ± 2.97%) of migratory Gfy traced neurons in the nose were labelled with EdU at these stages (Fig. S5). Altogether, this suggests that that during embryonic development GfyCre-mediated recombination is variably detected between 2 and 4 days after neurogenesis.

**Figure 3.**
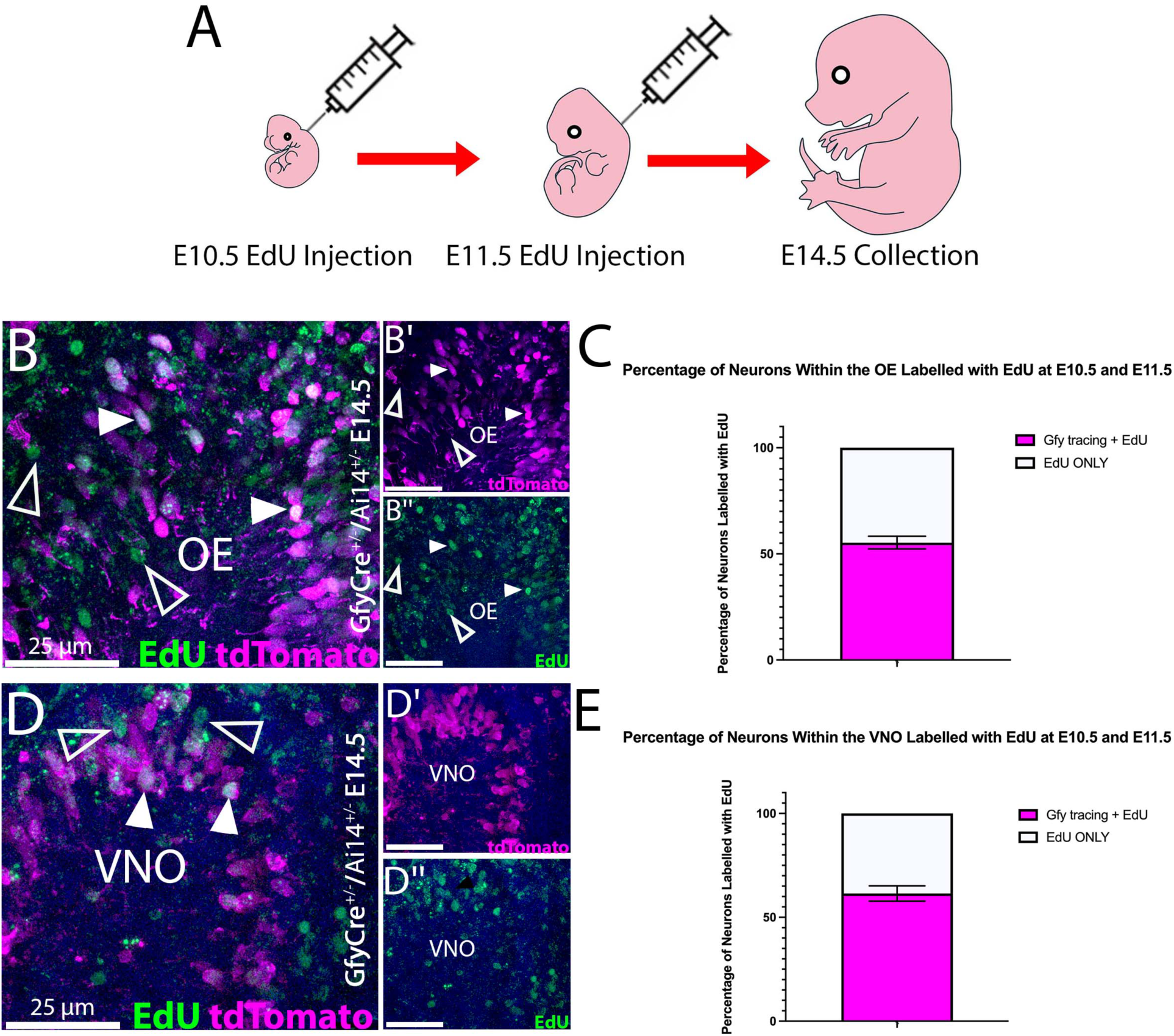
Gfy lineage tracing labels early-born chemosensory neurons. A) Schematic of EdU birth-dating experiments: EdU injections at E10.5 and E11.5 followed by tissue collection at E14.5. B–B″) GfyCre^+/−^;Ai14^+/−^ E14.5 olfactory epithelium (OE) sections stained for EdU (green) and Gfy lineage tracing (tdTomato, magenta). Gfy-traced neurons (open arrowheads) include a substantial fraction that is EdU-positive (filled arrowheads). C) Quantification of EdU-positive neurons in the OE showing the proportion that is Gfy-traced versus EdU-only (n=3). D–D″) GfyCre^+/−^;Ai14^+/−^ E14.5 vomeronasal organ (VNO) sections stained for EdU and Gfy tracing. Gfy-traced neurons (open arrowheads) showed a similar pattern to the OE within the VNO, with many that are EdU-positive (filled arrowheads). E) Quantification of EdU-positive neurons in the VNO showing the proportion that is Gfy-traced versus EdU-only (n=3). Scale bars B-D’’, 25 µm.

### During migration, Gfy is not expressed by the GnRH-1 neurons

After confirming that GfyCre mice exhibited the expected recombination in chemosensory neurons within 3 days from neurogenesis, we aimed to understand if the Gfy lineage included the early neurons forming from the placode such as the TN neurons, GnRH-1 neurons, or other neuronal populations that form in the nose and migrate to the brain (Hilal et al., 1996; Murakami et al., 2022; Murakami et al., 2024). Notably, recent RNA-sequencing data of the GnRH-1 neurons did not identify Gfy mRNA expression in migratory

GnRH-1 neurons (Zouaghi et al., 2025). The GnRH-1 neurons form between E10 and E12 (Jasoni et al., 2009; Schwanzel-Fukuda et al., 1989; Wray et al., 1989). We previously used Cre mouse lines to genetically label the GnRH, which indicates that these cells express the R26 reporter at detectable levels (Taroc et al., 2020a).

To trace different neurogenic waves of the GnRH-1 neurons in GfyCre/Ai14 embryos, some dams were injected with EdU at E8.5 and E9.5, and others at E10.5 and E11.5 (Fig. S6). We found that 15% (SD ± 5.35%) of the GnRH-1 neurons were labeled by EdU injection at E8.5 and E9.5 (Fig. S6B-C’’), while 40% (SD ± 7.59%) were labeled after injections at E10.5 and E11.5 (Fig. S6D-F’’).

RNAscope analysis for Gfy mRNA expression confirmed the lack of detectable Gfy RNA expression in migratory GnRH-1 neurons (Fig. S2F,G). These data confirmed no Gfy RNA expression in migratory GnRH neurons as determined by others afte FACS sorting (Zouaghi et al., 2025). Moreover, between E11.5 and E15.5 we could see GnRH-1 neurons migrating in association with Gfy traced migratory cells (Fig.4B-D) but we could not identify any Gfy tracing in GnRH-1 neurons in the nasal area or in the brain (Fig. 4), suggesting that Gfy is not expressed, or its expressed at undetectable levels, in newly formed or maturing GnRH-1 neurons even 6 days after neurogenesis.

**Figure 4.**
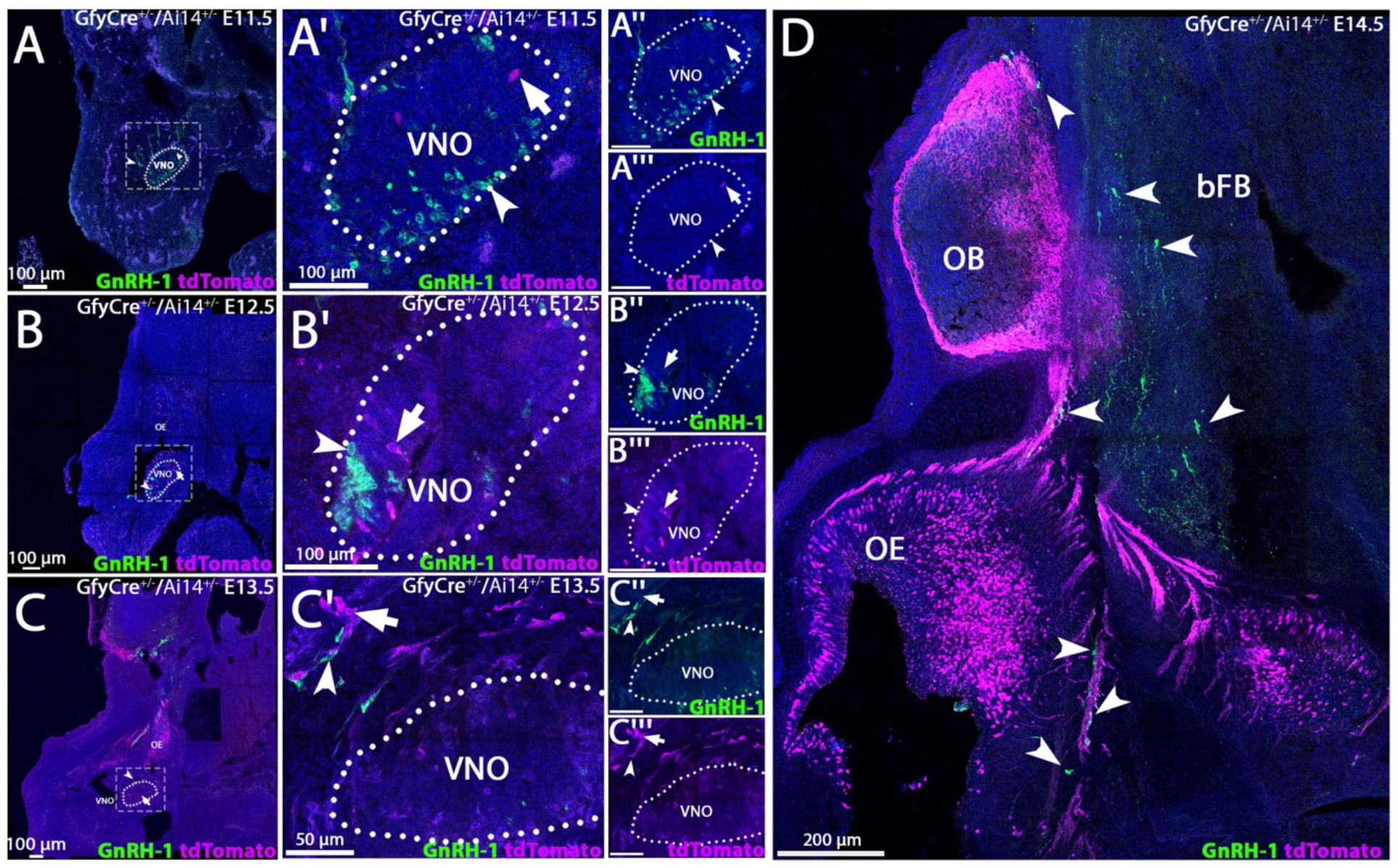
The GnRH-1 during their migration do not express Gfy. Parasagittal sections from GfyCre^+/-^;Ai14^+/-^ embryos at E11.5-E14.5 stained for GnRH-1 (green)(arrowheads). A-A‴) At E11.5, GnRH-1 neurons are located within and emerging from the vomeronasal organ (VNO), while Gfy lineage tracing (magenta) is sparse (arrows). B-B‴) At E12.5, Gfy-traced neurons increase in the VNO and olfactory epithelium (OE), coinciding with continued GnRH-1 neuronal migration. C-C‴) By E13.5, GnRH-1 neurons begin invading the brain, whereas Gfy tracing remains restricted to the nasal compartment. D) At E14.5, GnRH-1 neurons reach the basal forebrain (bFB); Gfy tracing is abundant in the nasal epithelium and detectable in the olfactory bulb (OB) but remains absent from GnRH-1 neurons. Scale bars: D, 200 µm; A-C and A’-B”’, 100 µm; C’-C”’, 50 µm.

### At completed migration, Gfy lineage-tracing highlights some GnRH-1expressing neurons as well as additional neurons in the preoptic area

To further test if Gfy expression and tracing could emerge at later developmental stages, after GnRH-1 migration has completed, we collected E18.5 embryos and postnatal animals at P56 (Fig. 5D-G), stages when the majority of GnRH-1 neurons have migrated to the brain (Wray, 2001). At E18.5, the GnRH-1 neurons could be identified in the brain close to the forebrain junction and scattered throughout the basal forebrain (Fig. 5A-A’’’) Analysis at E18.5 identified 3 populations of neurons in the basal forebrain: GnRH-1 neurons not traced, Gfy traced neurons negative for GnRH-1 immunostaining, and Gfy traced neurons positive for GnRH-1 immunostaining (Fig. 5A-C’’). From quantifications, we found that 12% (SD ± 1.12%; n=3) of GnRH-1 neurons in the brain at E18.5 were positive for Gfy tracing, a similar percentage to what we had previously reported after Prokr2Cre tracing (Amato et al., 2024). A rough count of Gfy traced neurons negative for GnRH-1 immunostaining established a average of 67 neurons (SD ± 11.9; n=3) At P56, GnRH-1 neurons were identifiable in the preoptic area (POA)/hypothalamic region (Fig. 5D-G). Notably, at this stage, we observed the same three neuronal populations as at E18.5, with similar representation as at E18.5, with 15% (SD ± 1.47%; n=3) of GnRH-1 immunopositive neurons in the POA traced for Gfy. Also, at this time, cells that were negative for GnRH-1 but positive for Gfy tracing could be found in the brain. Based on counts from 3 animals we estimated these to be 59 (SD ± 11.9; n=3) These findings indicate that contrary to what we observed during embryonic development, Gfy expression extends beyond the developing olfactory system, to additional migratory cells deriving that from the olfactory placode migrate in the brain.

**Figure 5.**
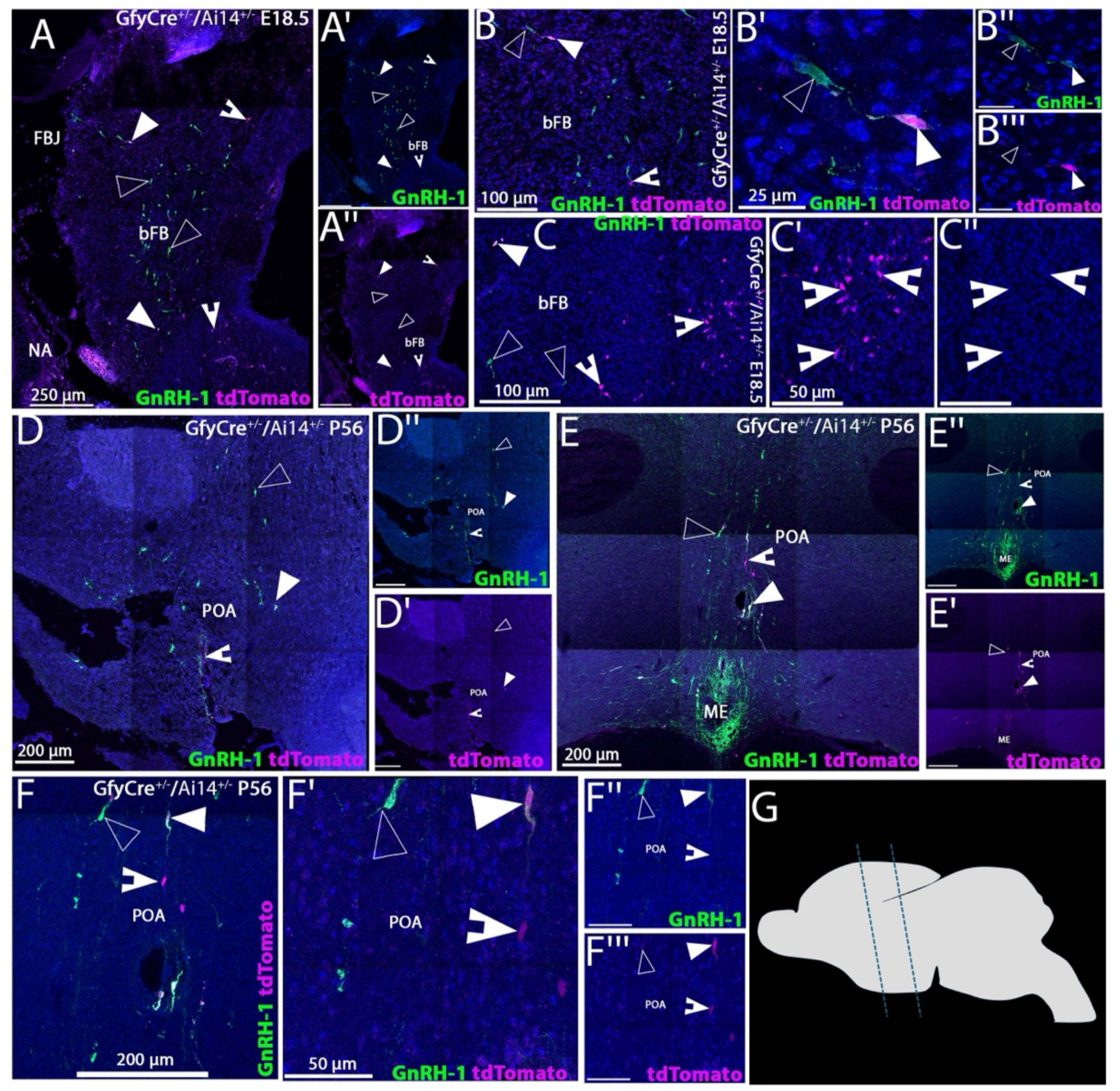
At completed migration, GfyCre-traced neurons are present in the basal forebrain. A-C’’) Immunoflourescent staining of GfyCre^+/-^;Ai14^+/-^ E18.5 embryos for GnRH-1 (green).GnRH-1 neurons are scattered from the point of entrance close to the forebrain junction (FBJ) to the basal forebrain (bFB), for reference use nasal area (NA). 3 staining patterns were observed: Gfy traced neurons (magenta, notched arrowheads), GnRH-1 neurons (empty arrowheads) and GnRH-1 neurons positive for Gfy tracing (white, arrowheads). D-F’’’) Coronal sections of P56 GfyCre^+/-^;Ai14^+/-^ including the preoptic area (POA) median eminence (ME), use G as reference for the brain areas analyzed. Gfy traced neurons (notched arrowheads) were observed proximal to the POA alongside GnRH-1 immunopositive Gfy traced neurons (arrowheads) and Gfy negative GnRH-1 neurons (empty arrowheads). G) Cartoon illustrating sagittal view of the brain, dotted lines delineate the areas of interest. Scale bars in A-A’’, 250 µm; D-D’’, E-E’’, F, 200 µm; B-B’’, C,100 µm; C’-C’’,F’-F’’’, 50 µm; B’-B’’’, 25 µm.

### Gfy is expressed by subsets of Prokr2 TN/pioneer neurons

In a recent paper, Prokr2 was identified as a reliable genetic marker for the neurons forming the TN as well as for putative olfactory pioneer neurons (Chen et al., 2026). In the developing nose, migratory cells positive for Prokr2Cre/Ai14 tracing can also be identified based on high immunoreactivity for Map2 up to E14.5 (Amato et al., 2024), notably the GnRH-1 neurons do not express Map2 (Herde et al., 2013). Analyzing GnRH-1 migration (Fig. 4D), we could see that the GnRH-1 neurons invading the brain at E14.5, were not associated with GfyCre traced fibers. In fact, the Peripherin+ fibers of the TN appeared negative for Gfy tracing (Fig. 2), though sparse axons positive for Gfy tracing could be found in the basal forebrain (Fig. S7B-B’’).

To determine whether Gfy-tracing could discriminate among different subpopulations of Map2–Prokr2–expressing migratory neurons and to exclude low detectability due to poor reporter expression among the TN fibers, we compared E14.5 GfyCre and Prokr2Cre lineage-traced embryos (Fig.6I-I’’).

**Figure 6.**
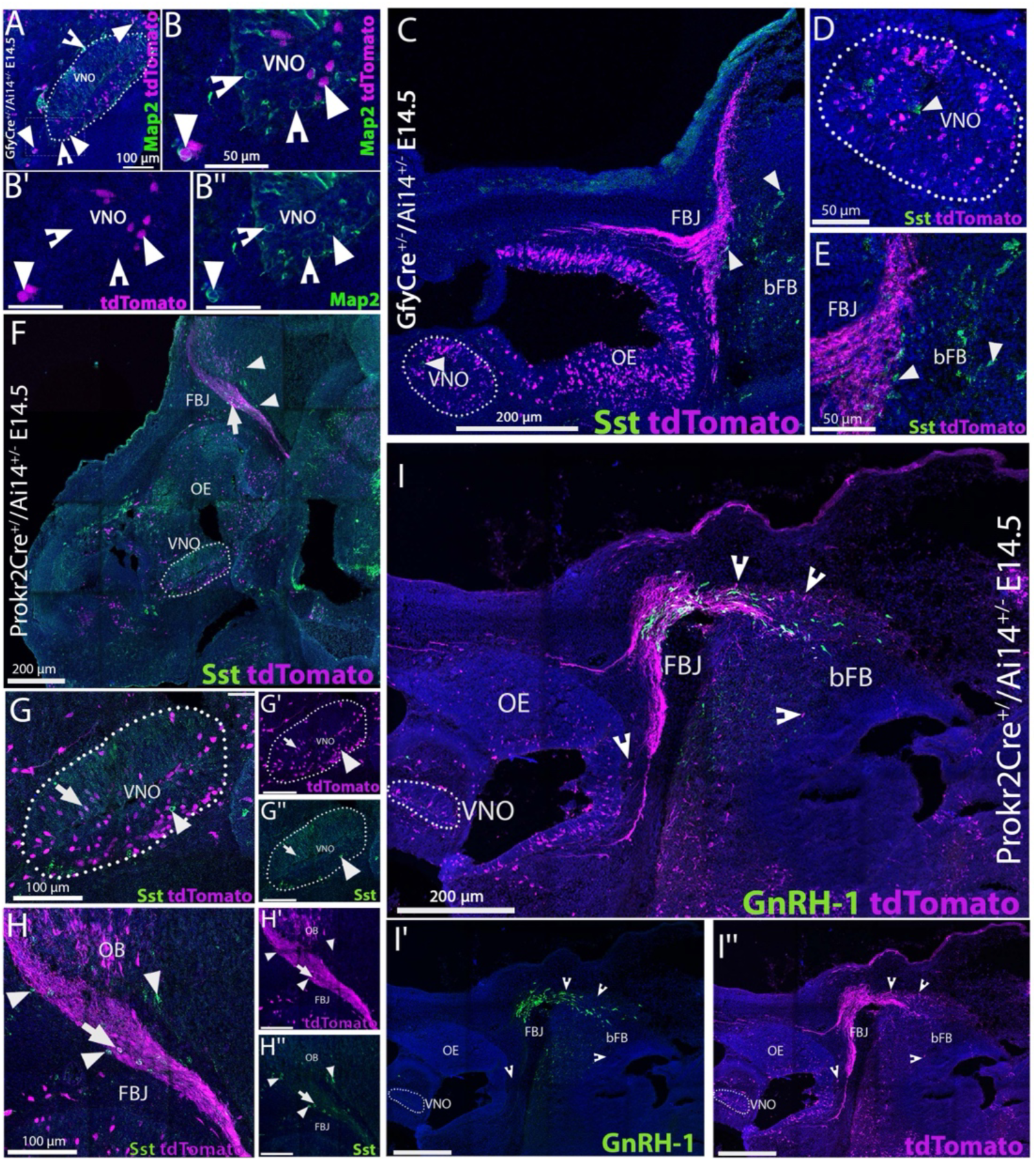
As GnRH-1 neurons migrate, Gfy traced migratory neurons are detectable outside of the brain. A-B’’) GfyCre^+/−^ /Ai14^+/−^ embryos at E14.5 paired with TN marker Map2 (green) immunofluorescent staining (notched arrows). Many Map2+ TN neurons were observed to migrate in clusters out of the vomeronasal organ (VNO). Gfy traced neurons (magenta) were observed in the VNO, and olfactory epithelium (OE) and migratory mass. Many of the migratory Gfy traced neurons appeared to be Map2+ (arrows). C-E) E14.5 GfyCre^+/−^ /Ai14^+/−^ embryos stained for Somatostatin (Sst, green). Sst neurons in the migratory mass near the forebrain junction (FBJ) and basal forebrain (bFB) are not positive for Gfy tracing (arrowheads). F-H’’) Prokr2Cre^+/−^ /Ai14^+/−^ E14.5 sections stained with Sst (green, arrowheads). Prokr2 tracing (magenta) appears to overlap with Sst expression (in both the VNO and migratory mass near the FBJ. I-I’’) Prokr2Cre^+/−^ /Ai14^+/−^ E14.5 embryos paired with GnRH-1 immunofluorescent staining (green). Prokr2 traced cell bodies and fibers migrate from the nose (notched arrowheads) invade the brain, distinct from the GnRH-1 neurons. Scale bars in B-B’’, D,E, 50 µm; A,G-G’’, H-H’’, 100 µm; C, F, I-I’’, 200 µm.

The TN/pioneer neurons were highlighted using Map2 as a marker. By pairing GfyCre tracing with Map2 immunostaining at E14.5, we found that some Map2-positive migratory neurons were positive for GfyCre tracing (Fig. 6A-B’’), suggesting that Gfy tracing highlights some TN/pioneer neurons. In line with this, anti-Gfy RNAscope on traced E14.5 Prokr2Cre traced embryos confirmed Gfy expression in traced TN-Pioneer neurons (Fig. S2H-I’’’).

Somatostatin (Sst) is a neuropeptide found to transiently expressed in numerous regions of the brain during embryonic development, with roles in neurite outgrowth and neuronal differentiation (Ferriero et al., 1994; Murakami and Arai, 1994; Song et al., 2021). Recent work in chick has found Sst expressing cells in the migratory mass (Murakami et al., 2022), we observed that also in mouse, cells expressing Sst at E14.5 emerge from the developing VNO and appear to migrate as part of the migratory mass toward the brain, however, Sst expressing cells were not positive for Gfy tracing (Fig. 6C-E). In contrast, by using Prokr2Cre tracing, we observed that Sst is expressed at least transiently by some of the Prokr2 traced migratory cells (Fig. 6F-H’’). In Prokr2 traced animals immunolabelled against Sst, we found that some of the Sst expressing cells were also positive for Prokr2 tracing in the migratory mass. However, Sst expressing cells in the brain appeared to be negative for both GfyCre and Prokr2Cre tracing at E14.5. (Fig. 6E, H-H’’), this was true also in postnatal animals (data not shown).

### Following Gfy expression in postnatal VSNs suggests subtle diCerences in the developmental dynamics of V1R and V2R VSNs

The expression of Gfy in newly generated neurons makes this line a valuable tool for manipulating chemosensory neurons prior to their maturation. Because of our focus on neuronal populations of and arising from the vomeronasal primordium, we further examined Gfy expression in this area. During embryonic development, we found that GfyCre recombined in both Meis2 and AP-2ε (Tfap2e) expressing neurons, which later will mature into V1R and V2R VSNs respectively (Fig. S8) (Enomoto et al., 2011; Katreddi et al., 2022).

To characterize GfyCre recombination in relation to postnatal vomeronasal neurogenesis, we reanalyzed whole-VNO single cell RNA-sequencing datasets (Hills et al., 2024; LeFever et al., 2024) and reconstructed the canonical dichotomy of apical and basal VSN development (Fig. 7A). UMAP projections and expression overlays for progenitors (Ascl1, Mki67, Neurog1), precursors (non-proliferative Neurog1, Neurod1), maturing VSNs (Gap43, Gng8/Gy8 and Gfy) and the maturation marker Omp. These recapitulated the expected neurogenic sequence, with progenitors at the base of the Y-trajectory, transitioning through neuronal precursors and immature VSNs, and diverging into the apical and basal branches, which culminate in mature V1R/Meis2+ and V2R/AP-2ε+ neurons (Fig. 7B). The transitional states of our single cell sequencing data was further confirmed through pseudotime reconstruction, recapitulating our classification of the cell clusters (Fig. 7H).

**Figure 7.**
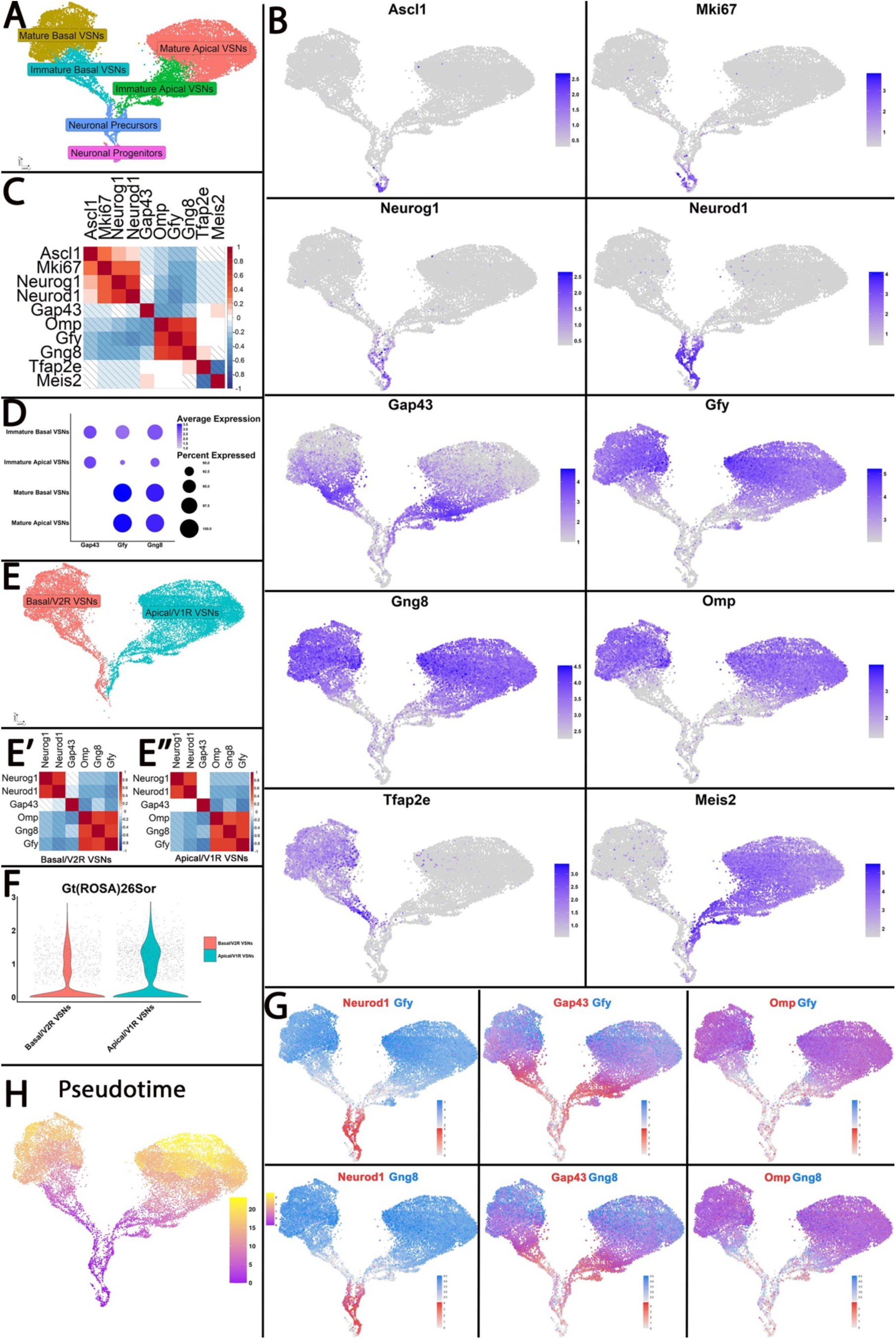
Single-cell transcriptomic analysis of the postnatal vomeronasal organ. A) UMAP projection of single cell RNA-sequencing data from the postnatal vomeronasal organ illustrating the major neuronal populations, including neuronal progenitors, neuronal precursors, immature apical and basal vomeronasal sensory neurons (VSNs), and mature V1R/Meis2+ and V2R/AP-2ε (Tfap2e)+ VSNs. The characteristic Y-shaped dichotomy of apical and basal lineages is resolved. B) UMAP feature plots showing expression of key developmental markers across the dataset. Ascl1 and Mki67 label proliferative progenitors, Neurog1 and Neurod1 label neuronal precursors, Gap43 marks immature neurons of both lineages, whereas Omp labels mature VSNs. Gfy and Gng8 are associated with early postmitotic neuronal states. AP-2ε and Meis2 highlight the basal and apical lineages, respectively. C) Correlation matrix illustrating co-expression patterns among canonical developmental and lineage-specific markers. Strong positive expression correlations identify the sequential transitions from progenitors to neuronal precursors to immature and mature VSNs. Negative correlation of expression indicated with diagonal lines. D) Dot plot summarizing Gap43, Gfy, and Gng8 expression across immature and mature apical and basal VSNs. Average expression and percent of expressing cells show that Gfy and Gng8 are lowly expressed in immature stages and are enriched upon maturation. E) UMAP projections showing the apical (V1R/Meis2+) and basal (V2R/Tfap2e+) VSN lineages segregated for comparison. E′–E″) Correlation matrices demonstrating that the developmental progression of apical and basal VSNs involves similar transcriptional transitions, but with distinct lineage-specific marker relationships. F) Violin plots comparing expression of the Gt(ROSA)26Sor locus between basal and apical VSNs. (G) Co-expression UMAPs illustrating the spatial and developmental relationships of Gfy and Gng8 with Neurod1, Gap43, and Omp, highlighting that Gfy and Gng8 mark neurons before and after they reach full maturation. H) Pseudotime reconstructions show early pseudotime (purple) and late pseudotime (yellow), this shows the developmental progression of cell populations present in the VNO.

Pearson correlation plots were generated to represent the linear relationship between two genes using the Pearson correlation coefficient, ranging from positive correlation (+1), negative correlation (-1) and no correlation (0). Correlation plots revealed consistent transcriptional changes across the dataset (Fig. 7C). This analysis showed that in contrast to what was observed during embryonic development (Fig. 1J), a negative correlation between Gfy and Gap43, along with a positive correlation between Gng8 and Omp. This suggests that postnatally, Gfy is expressed with a delay relative to Gap43 in maturing neurons and maintains high expression in mature neurons, while Gng8 clusters with intermediate neuronal states (Fig. 7D,G).

When the apical and basal branches were examined separately, we found additional interesting differences in their internal development (Fig. 7E–E″). In fact, we noticed that Gap43 expression rises earlier in Meis2+/apical lineage as opposed to AP-2ε+/basal lineage (Fig. 7E’, E’’). On the contrary, the Meis2-/basal lineage was found to have a negative correlation between Neurod1 and Gap43.

Further analysis at precursor stages reinforced this difference (Fig. S9), showing higher Gap43 and Gfy expression in the Meis2+/apical lineage. Moreover, by performing correlation analysis on Meis2+/apical and AP-2ε+/basal precursor populations, we observed that Meis2+ cells have a more precocious expression of Gfy, partial overlapping with Neurod1 and Omp (Fig. S9C’-C’’). Suggestive of differences in developmental dynamics between the two main types of VSNs.

From previous work we know that V2R VSN specific transcription factor AP-2ε directly binds to the Gt(ROSA)26Sor locus, potentially modulating its expression in V2R neurons (Lin et al., 2022). Based on that, we assessed expression of the Gt(ROSA)26Sor locus to evaluate baseline reporter activity across VSN classes. Violin plots revealed a modest but consistent elevation in ROSA expression in apical (Meis2+) VSNs relative to basal (AP-2ε+) neurons (Fig. 7F), suggesting lineage-specific differences that may influence reporter sensitivity in genetic tracing experiments.

To validate some of these observations in vivo, we combined GfyCre lineage tracing with Gap43 immunofluorescence (Fig. 8A-A’’’). At P30, GfyCre-traced neurons were present in both immature and mature VSNs. However, a difference in the intensity of tdTomato expression was visibly noticeable between neurons in the apical and basal territories (Fig 8A-A”).

**Figure 8.**
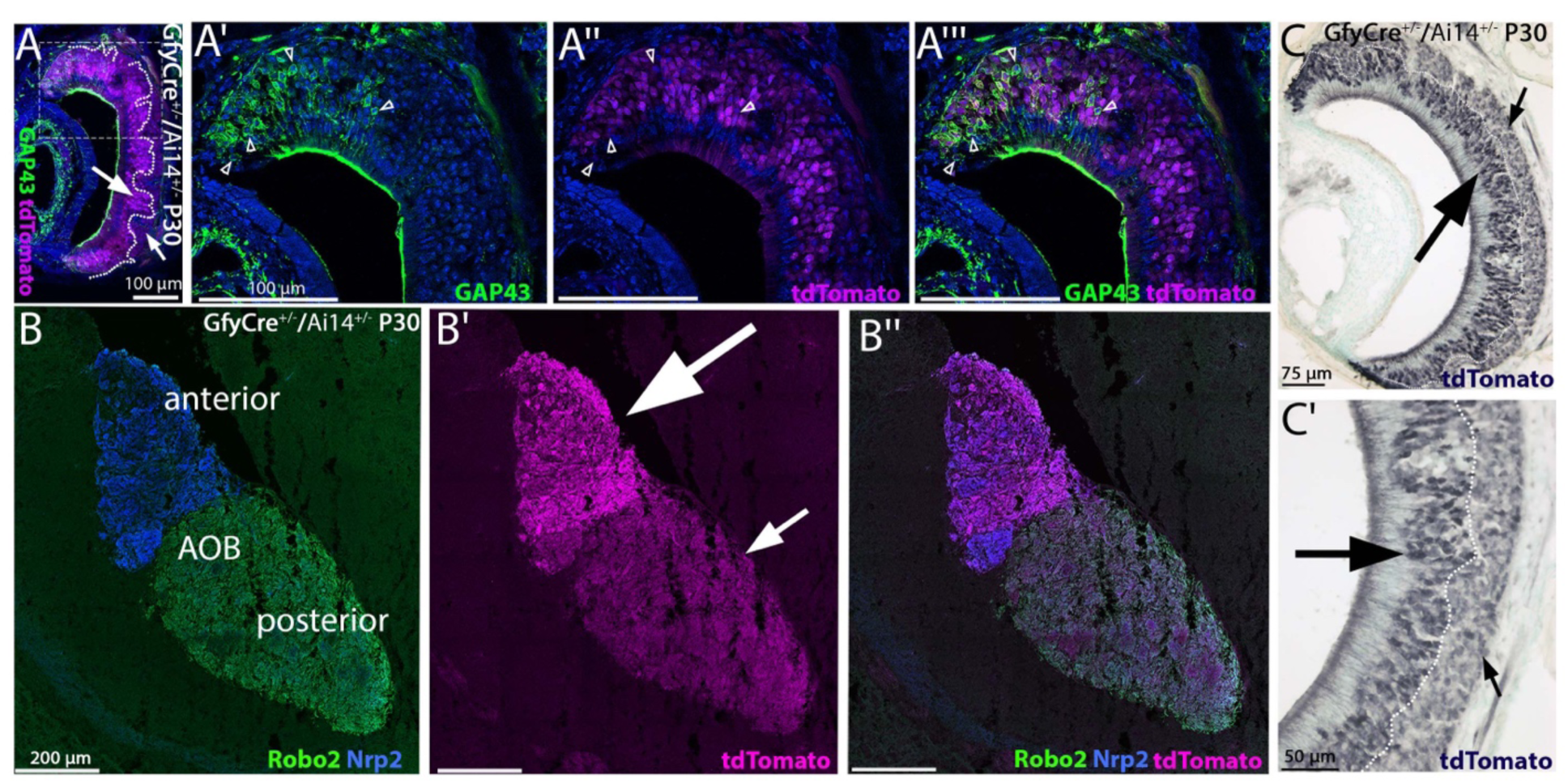
GfyCre-traced neurons in the VNO and AOB exhibit diCerent levels of ROSA26 reporter expression. A) Postnatal day 30 (P30) GfyCre^+/-^;Ai14^+/-^ vomeronasal organ (VNO) coronal sections stained for the immature vomeronasal sensory neuron (VSN) marker Gap43 (green) and tdTomato (magenta). Gfy lineage tracing is detected at lower levels in neurons in the basal territories (small arrows) and at higher levels in VSNs in the apical regions (larger arrows). A’-A”’) enlargements of the marginal zone (boxed in (A)) showing GAP43; tdTomato and merge A”’). Arrows indicate GAP43 (green) immunoreactive cells negative for tdTomato or expressing below detectability. B–B″) P30 GfyCre^+/-^;Ai14^+/-^accessory olfactory bulb (AOB) sections stained for Robo2 (green) and Nrp2 (blue) to distinguish posterior and anterior AOB projections, respectively. GfyCre tracing (magenta) is present in both anterior and posterior AOB regions but appears weaker in the Nrp2-positive posterior AOB (small arrow) compared with the anterior AOB (large arrow). C-C’) Nickel DAB intensified (black) immunohistochemistry of tdTomato on P30 GfyCre^+/-^;Ai14^+/-^ coronal VNO sections. TdTomato tracing signal in the apical region appears darker (large arrows) than in the basal region (small arrows). Scale bars: C’, 50 µm; C, 75 µm; A–A‴, 100 µm; B–B″, 200 µm.

As expected from their known axonal guidance programs, V1R/Meis2+ VSNs (Neuropilin-2+) projected to the anterior AOB, whereas V2R/AP-2ε+ VSNs (Robo2+) projected to the posterior AOB. GfyCre-mediated recombination was detected in the axons targeting both AOB domains (Fig. 8B-B’’). The reduced Rosa26 expression characteristic of basal VSNs (Fig. 8C-C’) again translated into visibly weaker tdTomato signals in the posterior AOB glomeruli compared to those in the anterior AOB glomeruli (Fig. 7F).

## Discussion

The lineage and developmental relationship between olfactory sensory neurons (OSNs) and neurons of the migratory mass in the developing nose has long been debated. Early observations showed that GnRH-1 neurons emerge from the olfactory placode (OP) and migrate alongside olfactory axons toward the forebrain, suggesting that a branch of the vomeronasal nerve might provide cues guiding GnRH-1 neurons to the basal forebrain (Hanchate et al., 2012; Schwanzel-Fukuda et al., 1989; Wierman et al., 2011; Wray et al., 1989). This hypothesis was reinforced by studies of Kallmann syndrome, in which anosmia often co-occurs with failed GnRH-1 neuronal migration, implying a shared developmental dependency (Kallmann FJ, 1944; Schwarting et al., 2001; Yoshida et al., 1995). However, subsequent studies suggest that the connection between OSNs and GnRH-1 neurons is less direct than originally thought. Increasing evidence including the ones reported here, indicates that the TN neurons, a neuronal population different from OSNs and VSNs, is responsible for guiding the GnRH-1 neurons into the brain (Amato et al., 2025a; Amato et al., 2024; Murakami et al., 2024; Taroc et al., 2019; Taroc et al., 2020b; Taroc et al., 2017). This view has shifted the field toward a model in which multiple placode-derived neuronal populations contribute specialized roles in GnRH-1 migration (Murakami et al., 2022).

Here we characterized neuronal populations in the developing olfactory system using the Gfy/123Cre lineage-tracing line (Takeuchi et al., 2010). Approximately 50% of GfyCre animals showed early recombination preceding Gfy expression in the nasal compartment, likely reflecting developmental mosaic recombination (Fig. S1). Because of this variability, maintaining this line on a reporter background is important to monitor recombination patterns (Fig. S1D). The remaining animals showed the expected recombination restricted to nasal chemosensory neurons.

Single-cell RNA-seq analysis at E14.5 revealed that Gfy transcripts are enriched in differentiated olfactory neurons as well as subsets of migratory neurons within the nasal compartment (Fig. 1). Gfy expression largely overlaps with Gap43 during embryonic development, consistent with expression in postmitotic neurons. In vivo lineage tracing confirmed these observations: GfyCre;Ai14 embryos showed progressively increasing recombination between E11.0 and E15.5 in OSNs, VSNs, and Grueneberg ganglion neurons, along with their projections to the developing olfactory bulbs (Fig. 2, S3). RNAscope further identified Gfy-expressing neurons migrating away from the chemosensory epithelia toward the developing olfactory bulbs (Fig. S2B–C). Together, these findings support previous reports linking Gfy expression to neuronal populations derived from the OP or associated chemosensory epithelia (Baker and Farbman, 1993; Habif et al., 2023; Hirata et al., 2006; Kaneko-Goto et al., 2013; Valverde et al., 1993). In contrast, GnRH-1 neurons showed no detectable Gfy expression during their migratory phase. RNAscope and lineage tracing revealed no overlap between GnRH-1 neurons and GfyCre recombination between E11.5 and E14.5 (Fig. 4, S2F,G) which is in line with recent RNA seq data (Zouaghi et al., 2025).

RNA scope confirmed detectable Gfy expression as early as E11.5 (Fig S2A-C’’). Birth-dating experiments further confirmed that Cre recombination occurs in olfactory and vomeronasal sensory neurons approximately 3–4 days after neurogenesis, whereas migrating GnRH-1 neurons remain negative for recombination even up to five days after their birth (Fig. 3). Nevertheless, after migration is completed, a small subset (12–15%) of GnRH-1 expressing neurons resulted positive for Gfy tracing in the brain. This was observed on samples at E18.5 and P56 (Fig. 5). This could reflect either transient low-level Gfy expression in GnRH-1 neurons, sporadic recombination, or the existence of distinct GnRH-expressing neuronal populations with different transcriptional programs. Based on previous studies we tend to exclude that lack of recombination across GnRH-1 and TN neurons results from low Rosa reporter expression in the migratory neurons (Amato et al., 2024; Taroc et al., 2020a). Notably, while Prork2 tracing labels a large portion of pioneer/TN migratory neurons GfyCre tracing only labels subsets of these cells (Amato et al., 2024) (Fig. S2H-I’’’, 6).

Our data therefore suggest that Prokr2Cre marks a broader population of migratory neurons, whereas GfyCre recombination is mostly restricted to neurons that remain in the nasal compartment. In agreement with this model, peripherin-positive TN fibers entering the basal forebrain together with GnRH-1 neurons were mostly negative for Gfy tracing (Fig. 2, S7). Both RNAscope analysis and previous RNA-seq datasets confirm that Gfy transcripts are absent in GnRH-1 neurons as they migrate to the brain (Zouaghi et al., 2025) (Fig. S2F-G).

Interestingly, Gfy lineage tracing highlights some cells in the basal forebrain of animals at late developmental stages (E18.5) and postnatal stages (P56), only after GnRH-1 neurons completed migration (Fig. 5). At E18.5 and in the postnatal hypothalamus, three populations could be distinguished: GnRH-1 neurons negative for Gfy tracing (∼85%), GnRH-1 neurons positive for Gfy tracing (∼15%), and Gfy-traced cells that do not express GnRH-1.

To interpret the outcome of our study, some technical considerations should also be noted. Low or transient Cre expression can reduce recombination efficiency and obscure lineage detection in specific neuronal populations. In line with this, we see that Gfy expression which drives Cre expression is lower and less penetrant in migratory neurons compared to chemosensory neurons (Fig. 1G, K). Future studies could address this limitation using the recently generated Gfy-Venus-IRES-Cre knock-in line, which allows simultaneous visualization of endogenous Gfy expression and Cre-mediated recombination (Bao et al., 2025; Enomoto et al., 2019). Such tools will enable more precise evaluation of lineage relationships and expression dynamics in developing nasal neuronal populations.

In addition to the embryonic analysis, the analysis of vomeronasal sensory neurons revealed that GfyCre recombination occurs in both apical (Meis2/V1R) and basal (AP-2ε/V2R) lineages, indicating that Gfy expression is not lineage-restricted (Fig. 7). However, transcriptomic analysis revealed differences in maturation dynamics between these populations. Apical VSNs show earlier and stronger overlap between Gap43 and Gfy expression, whereas basal VSNs display delayed or weaker expression patterns (Fig. 7E–E’’) (Hills et al., 2024; Lin et al., 2018). These intrinsic differences extend to reporter sensitivity at the Rosa26 locus, which shows lower baseline expression in basal VSNs and likely contributes to weaker lineage-tracing signals in this population (Lin et al., 2022). In vivo validation confirmed that GfyCre recombination occurs after early differentiation markers such as Gap43 but robustly labels neurons as they mature and project to the accessory olfactory bulb (Fig. 8).

Together, these findings establish Gfy expression as a shared molecular feature across various postmitotic neurons forming in the OP. Notably, the differential levels and dynamic expression can be used to further refine the interpretation of lineage tracing in the olfactory system.

Importantly, our results, revealing either low or no Gfy expression, highlight additional genetic differences between GnRH-1 neurons and TN populations from canonical OSN and VSN lineages, supporting a model in which multiple placode-derived neuronal populations contribute to the formation of the migratory scaffold guiding GnRH-1 neurons into the brain.

## Methods

### Animals

Gfy(123)-Cre mice were provided generously from Dr. Thomas Bozza (Northwestern University), originally generated at RIKEN Brain Science Institute, Japan (Dr. Yoshihiro Yoshihara). PCR analysis was performed for genotyping, using generic Cre primers as follows: Cre FWD, AGG TGT AGA GAA GGC ACT TAG C, *Cre RVS, CTA ATC GCC ATC TTC CAG CAG G.* Ai14 mice were genotyped following the B6.Cg-*Gt(ROSA)26Sor^tm14(CAG-tdTomato)Hze^*/J protocols available on jax.org. Generation of Prokr2Cre^+/−^ mice (Mohsen et al. **<u>2017</u>**) and genotyping (Amato Jr. et al. **<u>2024</u>**) were previously described. CO_2_ euthanasia was performed, then immediate cervical dislocation. Mice of either sex were used for experiments. All animal experiments and procedures were completed in accordance with the guidelines of the Animal Care and Use Committee at the University at Albany, SUNY.

## Tissue Preparation

### Embryonic Tissue Collection

Embryo collection was carefully performed on time-mated females, where observation of the vaginal plug was estimated to be embryonic day 0.5 (E0.5). Collected embryos were placed into 3.7% formaldehyde/PBS at 4°C for 2 h to fix. Embryos then were submerged in 30% sucrose overnight, then embedded and frozen in O.C.T (Tissue-Tek) before storing at −80°C. The CM3050S Leica cryostat was used to cryosection embedded embryos, tissue sections were collected using Superfrost plus slides (VWR) at 18 µm thickness for immunofluorescent staining.

### Postnatal Tissue Collection

P30 and P56 mice were sedated with Sodium Pentobarbital, then transcardial perfusion was performed with 1X PBS followed by 4% Paraformaldehyde in PBS. Noses were separated from brains, with the noses being post fixed in 4% Paraformaldehyde overnight at 4°C.

Brains were post fixed in 1% Paraformaldehyde overnight at 4°C. The following day, brains were cryoprotected with 30% Sucrose in 1X PBS at 4°C overnight and noses were incubated in 500mM EDTA in 1X PBS for decalcification. At end of EDTA incubation noses were cryoprotected in sucrose. Samples were embedded and frozen in O.C.T and stored in -80°C prior to cryosectioning serial sections on Superfrost plus (VWR) slides. Both P30 noses and P56 brains were sectioned in a coronal orientation, while P30 brains were sectioned in the parasagittal orientation.

### Immunofluorescence

The following primary antibodies and dilutions were used for this study: goat-α-RFP (1:1000, Rockland), chicken-α-Peripherin (1:500, Abcam), mouse-α-HuCD (Invitrogen, 1:200), SW rabbit-α-GnRH-1 (1:6000, Susan Wray, NIH), rabbit-α-Map2 (1:150, Cell Signaling Technology), mouse-α-RFP (1:500, Origene), rabbit-α-Somatostatin (1:100, Abcam), rabbit-α-Gap43 (1:1000, Abcam), goat-α Neuropilin2 (1:4000, R&D Systems), mouse-α-Robo2 (1:50, Santa Cruz), rabbit-α-RFP (1:500, Rockland), goat-α-OMP (1:4000, WAKO), goat-α-AP-2ε (1:500, R&D Systems), and rabbit-α-Meis2 (1:1000, Abcam).

Antigen retrieval was used for experiments utilizing rabbit-α-Map2, mouse-α-RFP, goat-α Neuropilin2, mouse-α-Robo2, rabbit-α-RFP, goat-α-AP-2ε, and rabbit-α-Meis2, slides were submerged in a citric acid solution, gradually heated to 95°C, once at 95°C, boiling for 15 min. Slides were then cooled for 15 min before proceeding with standard immunohistochemical protocol.

For immunofluorescent staining, secondary antibodies selected for the correct species were conjugated with Alexa Fluor-488, Alexa Flour-594, or Alexa Flour-680 (Invitrogen and Jackson Laboratories). 4′,6′-diamidino-2-phenylindole (DAPI, 1:3000; Sigma-Aldrich) was used to counterstain sections and coverslips were mounted using Fluorogel l (Electron Microscopy Sciences). Confocal microscopy image acquisition was taken on a LSM 710 microscope (Zeiss). Epifluorescence images were acquired on a DM4000 B LED fluorescence microscope equipped with a Leica DFC310 FX camera. Whole embryo images were acquired using the Nikon SMZ25 stereo microscope. Image analysis and quantifications were done using FIJI/ImageJ software. Each staining presented was replicated on three different animals.

### Chromogen-Based Reactions

Chromogen-based reactions were performed as described previously (Forni, Fornaro, et al. 2011). Sections were processed using a standard avidin–biotin-horseradish peroxidase/3,3-diaminobenzidine (DAB) procedure. Nickel (II) sulfate heptahydrate (Sigma) was used to visualize and intensify the DAB reaction (black), then counterstained with methyl green. Brightfield images were taken on a LeicaDM4000 B LED fluorescence microscope equipped with a Leica DFC310 FX 422 camera. Stainings presented were replicated on three different animals.

### Edu Administration and Detection

Edu was prepared according to ClickTech EdU Cell Proliferation Kit 488 for IM (Millipore-sigma, BCK-EDU488). Edu injections were performed on time-mated females previously described in the tissue preparation section of the methods. Intraperitoneal injections were done at various time points at a dosage of 50mg/kg body weight. We observed that development was not aNected by the injections. To detect Edu positive cells, we performed immunofluorescence as described in the previous section and followed the standard ClickTech EdU Cell Proliferation Kit 488 protocol. Slides were incubated for 30 minutes at room temperature in the prepared detection solution protected from light. After, we followed standard immunofluorescent protocol.

### RNAscope

Single-molecule fluorescence in situ hybridization coupled with immunofluorescence was performed using the RNAscope Multiplex Fluorescence v2 assay and the probe (RNAscope Probe-Mm-Gfy-C1 #1757711-C1) from ACDbio. RNAscope experiments were performed on 18 µm fixed-frozen E11.5-E14.5 mouse cryosections, following the manufacturer’s protocol.

### Cell Quantifications

Quantifications performed were done in FIJI/ImageJ using the cell counter plugin. All cell number quantifications in the nasal region were done on parasaggital E12.0-E18.5 tissue. For each quantification reported, cell bodies of neurons were counted. For all quantifications of cell counts, the number of cells per section (∼5 sections) was averaged for each animal.

### Embryonic Single-Cell Cell RNA Sequencing

Dimplots and feature plots were generated in R studio using the Seurat R package. The E14.5 single-cell sequencing data can be found on the Gene Expression Omnibus (GEO) database: GEO accession number GSE234871 (https://www.ncbi.nlm.nih.gov/geo/query/acc.cgi?acc=GSE234871). The data collection process and analysis was previously described (Amato Jr. et al. **<u>2024</u>**).

### Post-natal scRNA-seq quality control, clustering and analysis

#### Quality Control and Filtering

Postnatal single-cell RNA sequencing datasets (P14, P56, and P10, P21, P60) used for analysis can be found with GEO accession numbers <u>GSE247872</u>, and <u>GSE252365</u>. Data collection and analysis were previously described (Hills et al., 2024; LeFever et al., 2024). Independent ages respective datasets were merged and integrated to get a more full picture of the vomeronasal development. Cell filtering was done downstream to remove genes expressed in ≤3 cells, and ≥200 detectable genes. Quality control was done to clean up the data to only cells expressing between 200 and 9000 with ≤25% of those genes being mitochondrial genes.

#### CellCycle Regression

To avoid noise from cells undergoing active cell cycling, cells were scored for their features aligning with S, and G2/M phase as described in a SeuratV5 vignette (Hao et al., 2023). The difference between S.Score and G2M.Score was calculated, and these cells were regressed to include differences between cell types but remove differences within cycling cells.

#### DoubletFinder

The DoubletFinder program built for Seurat was applied to reduce and remove a predicted number of artificial doublets created during the process of single cell sequencing. This package was used as previously described (McGinnis et al., 2019).

#### Processing

Using SCTransform the data was scaled and normalized, the top 2000 highly variable features within the top 50 principal components were used for clustering. About 18,000 cells were used in this analysis and different developmental stages were identified through well documented key markers. When completing analysis each cluster/gene were compared against all other clusters/genes. In the Apical/V1R v Basal/V2R comparisons, only the Apical/V1R lineage was compared to the Basal/V2R lineage. Analysis of the differential Gt(ROSA)26Sor expression was conducted on P10, P21, P60 data previously published (LeFever et al., 2024). Pseudotime reconstruction was performed using the Mononcle3 R package.

## Acknowledgements

This work was supported by the Eunice Kennedy Shriver National Institute of Child Health and Human Development (NICHD) under Grants R01HD09733 (P.E.F.) and R01HD114827 (P.E.F.), as well as by the National Institute on Deafness and Other Communication Disorders (NIDCD) under Grant R01DC017149 (P.E.F.). The Zeiss 980 microscope at the University at Albany was funded by the Office of the Director, NIH, under Award Number S10OD028600.

We thank Dr. Yoshiro Yoshihara (Riken Center for Brain Science, Japan) and Dr. Thomas Bozza (Department of Neurobiology, Northwestern University, USA) for donating and sharing the Gfy Cre line with us.

## Supplementary Figures

**Figure S1.**
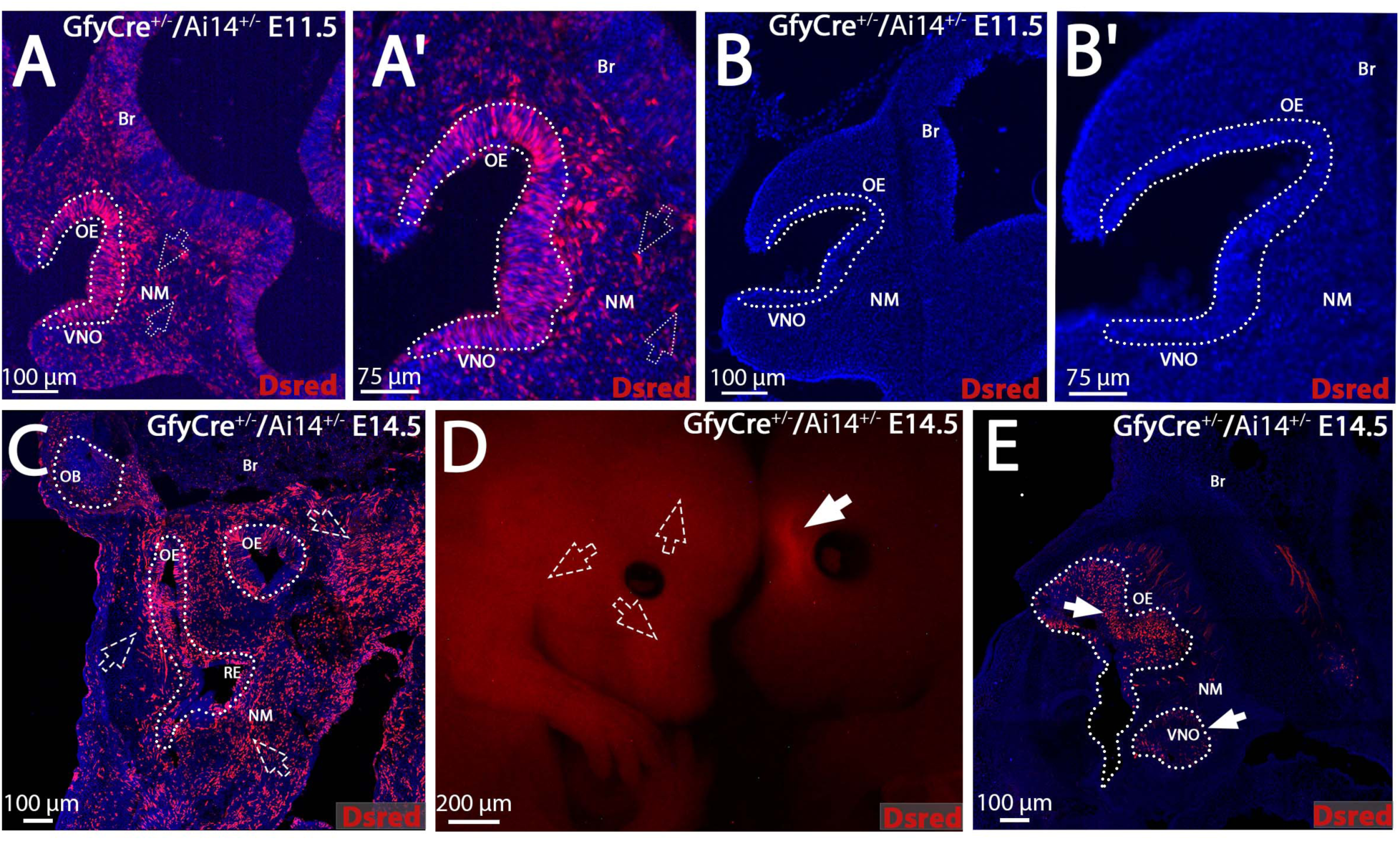
GfyCre Non-specific Recombination. A-B’) Both nonspecific and specific tracing patterns are observed in E11.5 GfyCre^+/−^/Ai14^+/−^ littermates. In the embryo showing nonspecific broad recombination (A-A’), the E11.5 cells positive for the recombination (red, empty dotted arrows) could be found in the nasal mesenchyme (NM), developing olfactory epithelium (OE), vomeronasal organ (VNO) and brain (Br); in contrast, the GfyCre^+/−^ /Ai14^+/−^ littermate does not show recombination at this stage. C) An E14.5 GfyCre^+/−^ /Ai14^+/−^ with nonspecific, broad chimeric tracing (red, empty dotted arrows) throughout the nasal mesenchyme (NM) the olfactory epithelium (OE), olfactory bulb (OB), respiratory epithelium (RE) and the brain (Br). D) Whole-embryo imaging comparing a GfyCre^+/−^ /Ai14^+/−^ embryo with nonspecific tracing present throughout the whole body (left, empty dotted arrows) and a GfyCre^+/−^ /Ai14^+/−^ embryo with correct tracing exclusive to the olfactory system (right, arrow). E) A traced GfyCre^+/−^ /Ai14^+/−^ E14.5 embryo, showing Gfy tracing (red, arrows) limited to the developing VNO and OE. Scale bars in D, 200 µm; A,B,C,D 100 µm; A’.B’, 75 µm.

**Figure S2.**
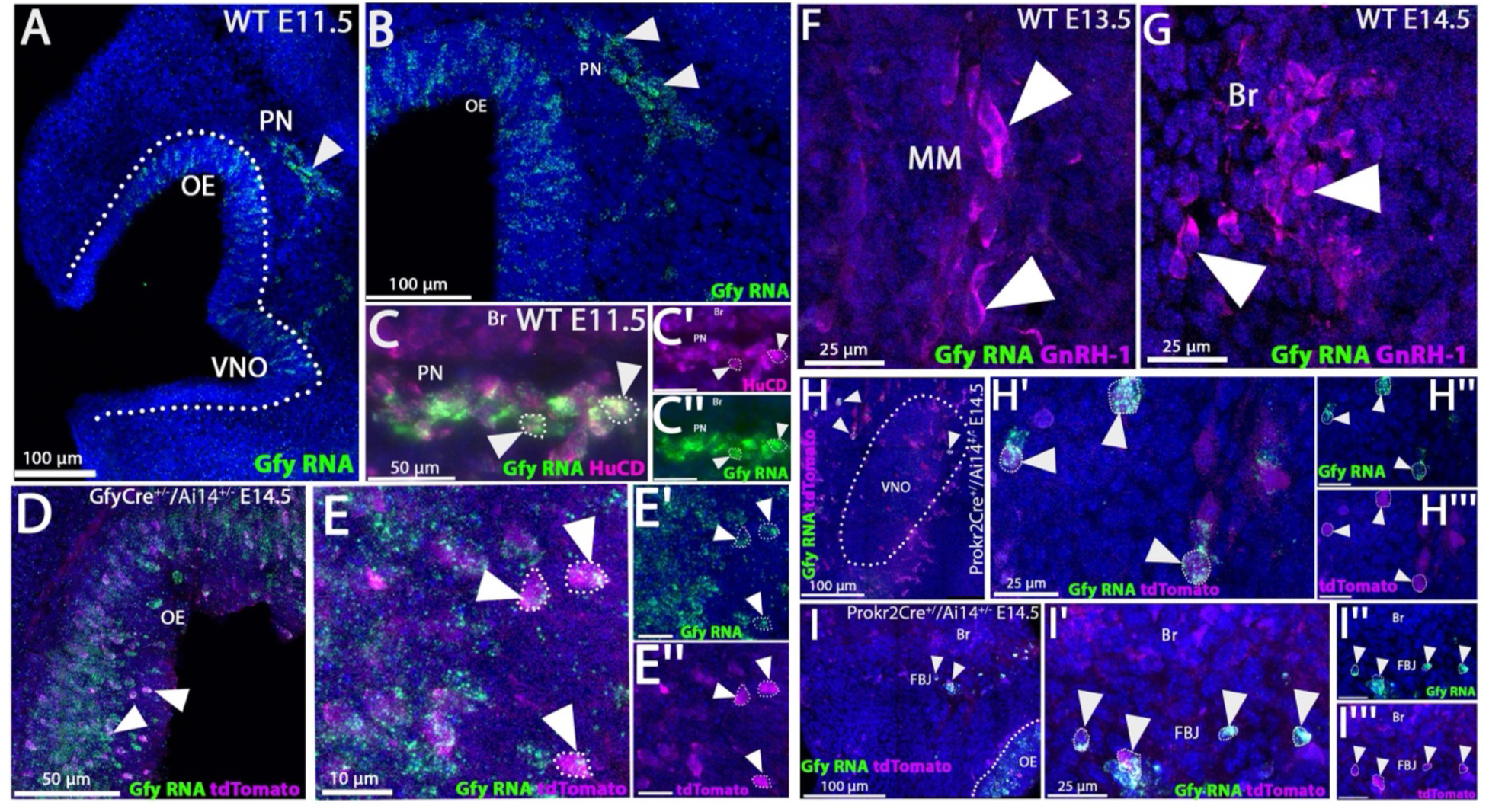
Gfy RNA expression in the developing olfactory system. A-B) Gfy RNAscope staining on E11.5 control animals, Gfy positive RNA puncta (green) were observed in the vomeronasal organ (VNO), olfactory epithelium (OE), and pioneer neurons (PN, arrows). C-C’’) Gfy RNA expression (green) paired with immunofluorescence for the neuronal marker HuCD (magenta), confirming that the Gfy RNA expression proximal to the brain (Br) are HuCD positive PNs (arrowheads, dotted outlines). D-E’’) GfyCre^+/−^ /Ai14^+/−^ E14.5 sections showed that Gfy traced neurons (magenta) were positive for Gfy RNA expression (green, arrowheads, dotted outlines). F-G) Gfy RNAscope paired with GnRH-1 immunofluorescent staining (magenta) on control E13.5 and E14.5 sections, GnRH-1 neurons in both the migratory mass (MM) and Br did not colocalize with any Gfy RNA expression (arrowheads). H-I’’’) Prokr2Cre^+/−^ /Ai14^+/−^ E14.5 sections showed Gfy expression in Prokr2 traced (magenta) terminal nerve/PNs (arrows) proximal to the VNO and forebrain junction (FBJ). Scale bars in A,B, H,I, 100 µm; C-C’’, D, 50 µm; F,G, H’-H’’’, I-I’’’, 25 µm; E-E’’, 10 µm.

**Figure S3.**
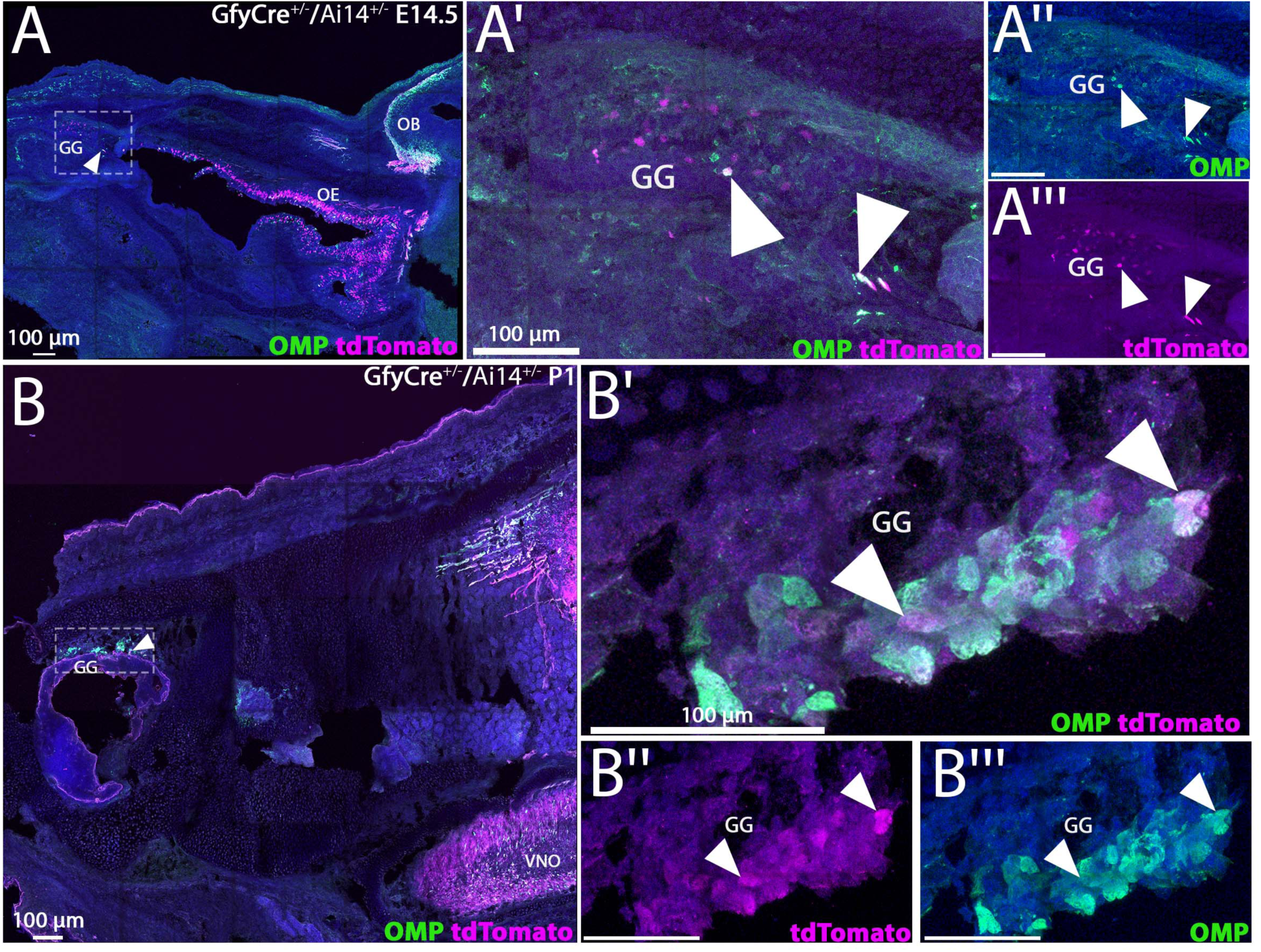
The Grueneberg Ganglion Expresses Gfy. A-A’’’) E14.5 GfyCre^+/−^ /Ai14^+/−^ embryos immunostained for Omp (green). Gfy tracing (magenta) can be observed in the olfactory bulb (OB), olfactory epithelium (OE) and overlapping with Omp expression in the neurons of the Grueneberg ganglion (GG) (white, arrowheads). B-B’’’) P2 GfyCre^+/−^ /Ai14^+/−^ animals show increased Gfy tracing in the vomeronasal organ (VNO) and GG. Scale bars in A-B’’’, 100 µm.

**Figure S4.**
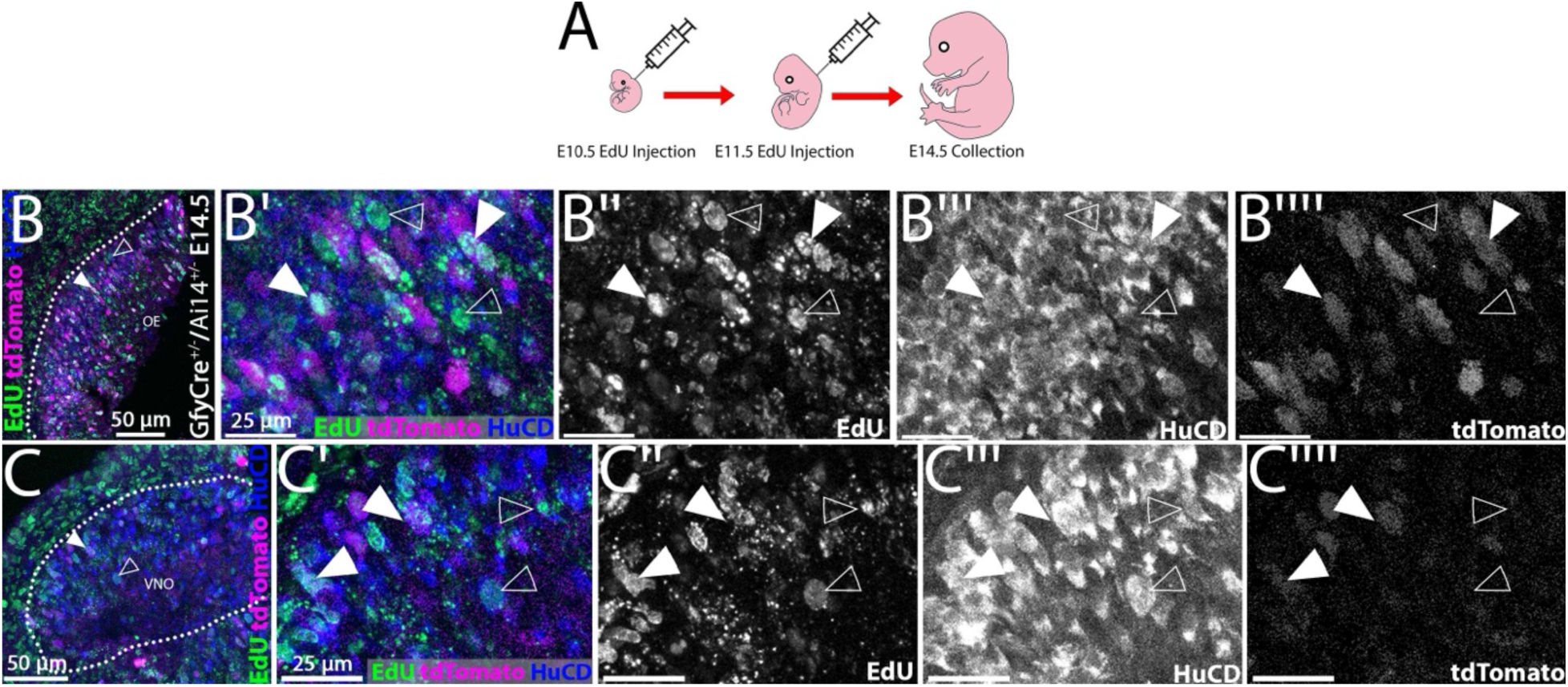
Gfy lineage tracing overlaps with HuCD expression and E10.5 and E11.5 EdU labelling. A) Schematic of E9.5 and E10.5 EdU labelling experiments, then collection at E14.5. B-C’’’’) Utilizing HuCD (blue) immunofluorescent staining, Gfy-tracing (magenta) and EdU staining (green), both Gfy tracing positive (arrowheads) and negative (empty arrowheads) neurons were observed to be labelled with EdU in the olfactory epithelium (OE) and vomeronasal organ (VNO). Scale bars in B, C, 50 µm; B’-B’’’’, C’-C’’’’, 25 µm.

**Figure S5.**
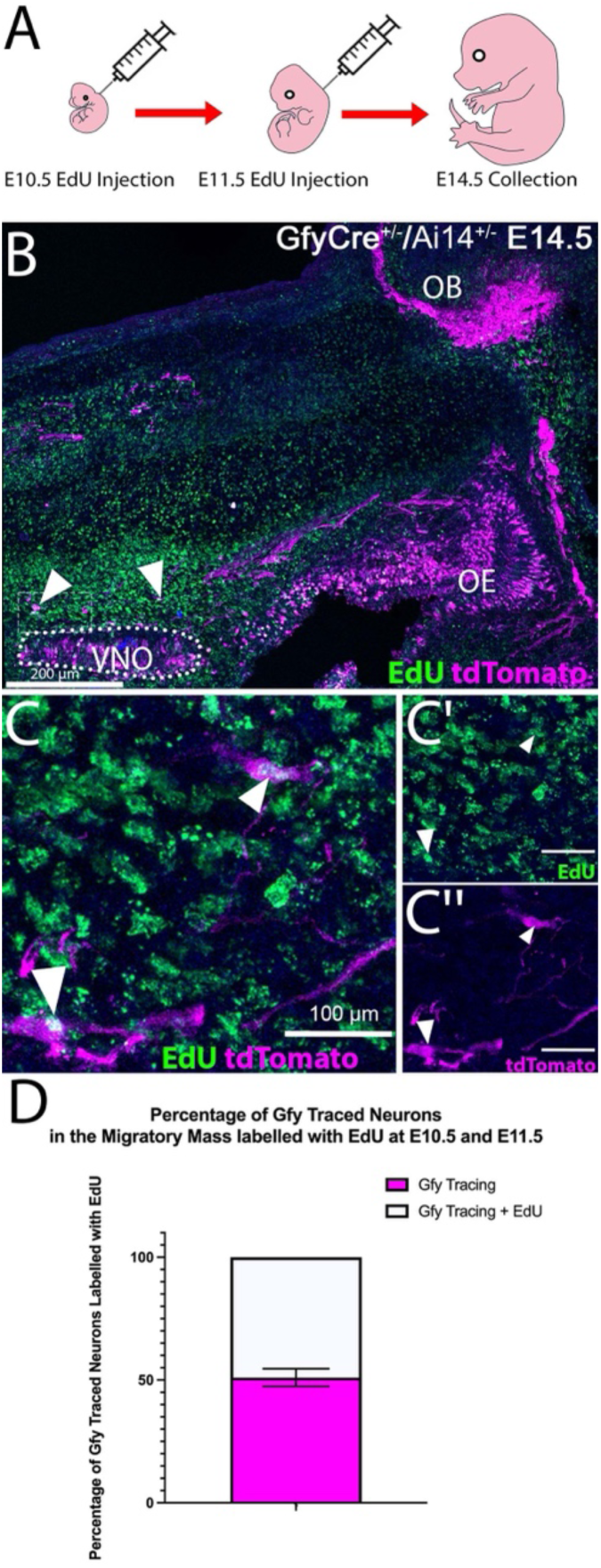
Gfy Tracing EdU Birth-dating Experiments in the Migratory Mass. A) Schematic of EdU injection experiments performed at E10.5 and E11.5 and subsequently collected at E14.5. B) Parasagittal section on E14.5 GfyCre^+/−^ /Ai14^+/−^ embryos stained for EdU (green) and Gfy tracing (magenta), with overlap highlighted by arrowheads. C-C’’) Gfy tracing (magenta) and EdU staining (green) overlap (white) indicated with the arrowheads. D) Quantifications of EdU positive Gfy traced migratory neurons in the migratory mass. Scale bars in B, 200 µm; C-C’’, 100 µm.

**Figure S6.**
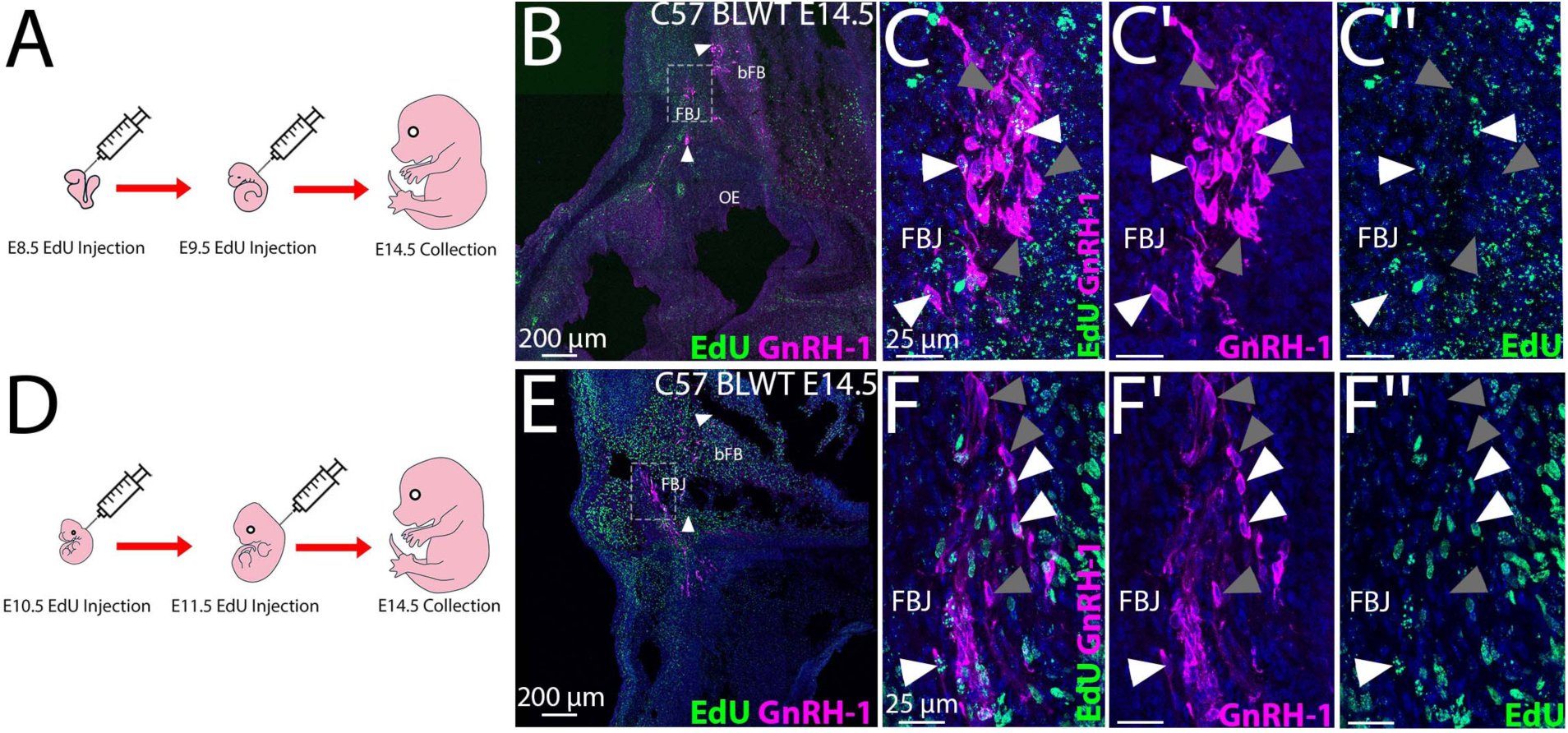
GnRH-1 EdU Birth-dating Experiments. A) Schematic of E8.5 and E9.5 EdU injection experiments, then collection at E14.5. B-C’’) E14.5 C57 BLWT control tissue immunostained for EdU (green) and GnRH-1 (magenta, grey arrowheads), overlap of EdU and GnRH-1 staining indicated with white arrowheads. D) Schematic of E10.5 and E11.5 EdU injection experiments, then collection at E14.5. E-F’’) EdU labelling at E10.5 and E11.5 was abundant in the forebrain junction (FBJ) and basal forebrain (bFB), overlapping with GnRH-1 immunoreactivity (white arrowheads). Scale bars in B, E 200 µm; C-C’’, F-F’’, 25 µm.

**Figure S7.**
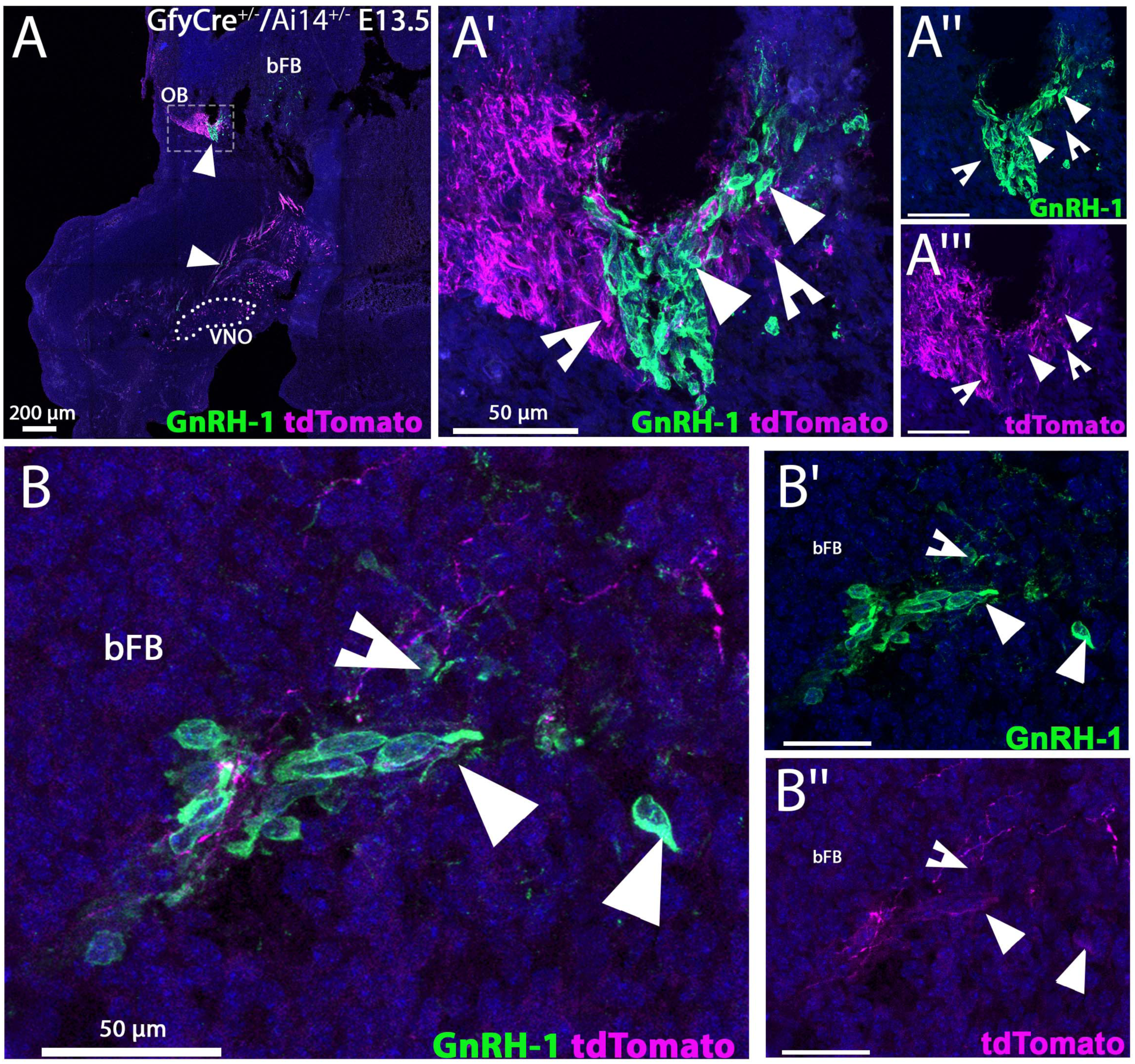
The GnRH-1 neurons are closely intertwined with Gfy traced fibers in the brain. A-A’’) E13.5 GfyCre^+/−^ /Ai14^+/−^ sections paired with GnRH-1 immunofluorescent staining. At this stage, GnRH-1 neurons (green, arrowheads) are migrating out of the vomeronasal organ (VNO), surrounded by Gfy traced cell bodies and axons/fibers (magenta, notched arrowheads) that also appear to be migrating out of the VNO. The GnRH-1 neurons invading the brain are not traced for Gfy. B-B’’) GnRH-1 neurons (arrowheads) invading the basal forebrain (bFB) with faint Gfy traced fibers (notched arrowheads) entangling the GnRH-1 neurons. Scale bars in A, 200 µm; B-B’’, C-C’’, 50 µm.

**Figure S8.**
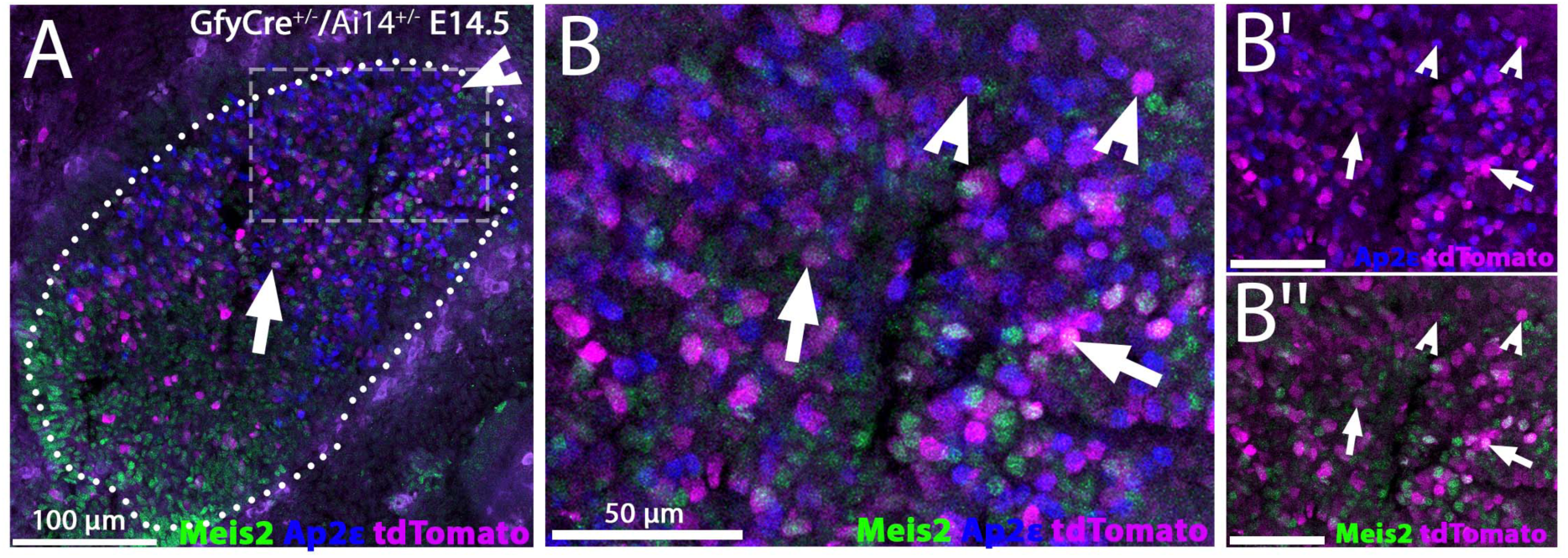
Genetic lineage tracing of Gfy in the embryonic VNO. A-B’’) E14.5 GfyCre^+/−^ /Ai14^+/−^ vomeronasal organ (VNO) sections stained for vomeronasal sensory neuron (VSN) populations, Meis2 (green, arrows, apical) and AP-2ε (blue, notched arrowheads, basal). Both apical and basal developing VSNs were positive for Gfy tracing at E14.5. Scale bars in A, 100 µm; B-B’’, 50 µm.

**Figure S9.**
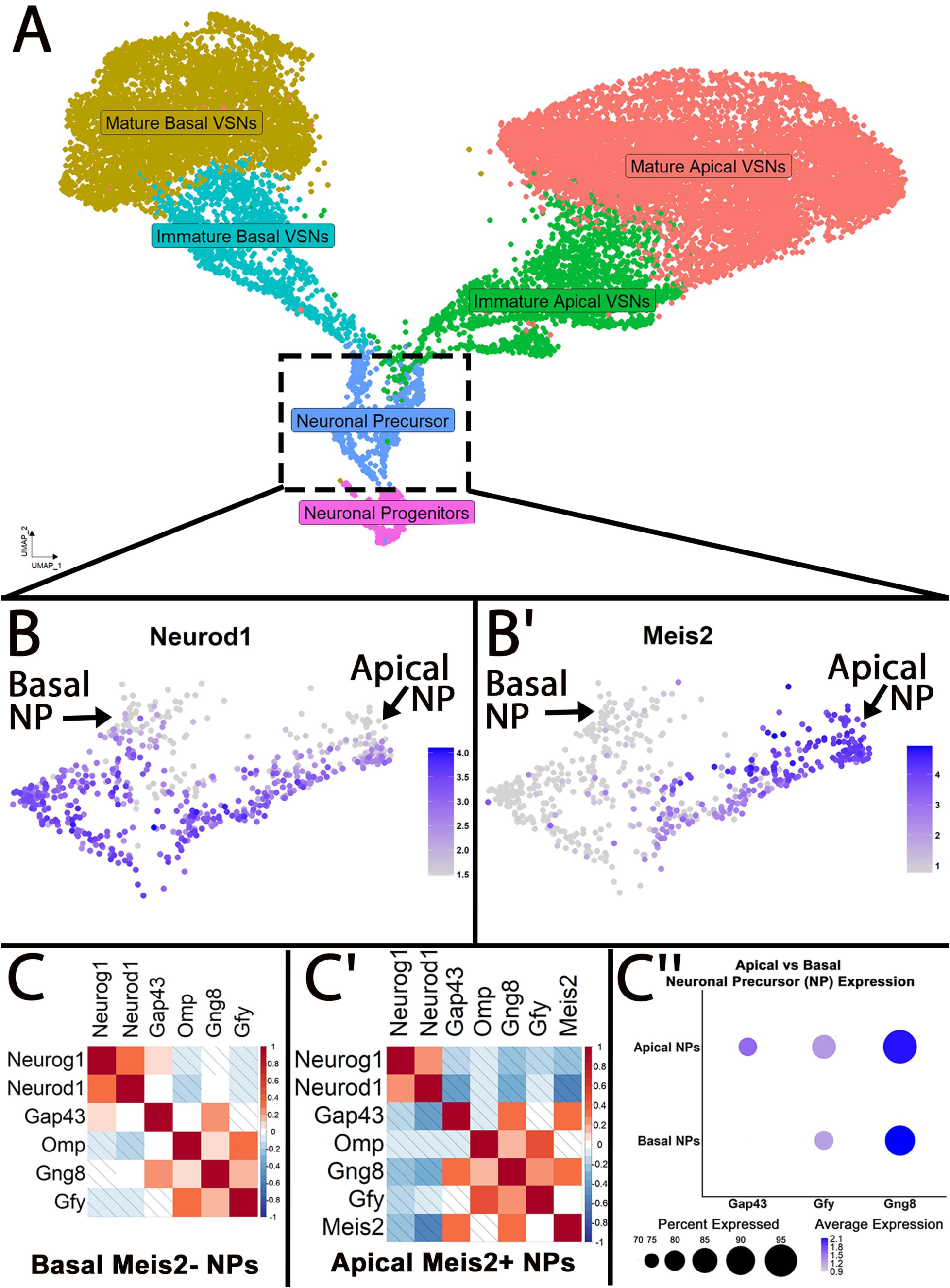
Single-cell transcriptomic analysis of the postnatal vomeronasal organ neuronal precursor populations. A) UMAP projection of scRNA-seq data from the postnatal vomeronasal organ illustrating the major neuronal populations: the neuronal progenitors, neuronal precursors (NP), immature apical and basal vomeronasal sensory neurons (VSNs), and mature V1R/Meis2+ and V2R/AP-2ε+ VSNs. B-B’) UMAP feature plots of the neuronal precursor population, Neurod1 highlights both apical and basal NP, while Meis2 expression is more enriched in apical neuronal precursors. C-C’) Correlation matrix showing gene expression correlations between the NP markers Neurod1 and key maturation markers (Gap43, Gng8, Gfy, Meis2). C’’) Dotplot illustrating the expression pattern of Gap43, Gfy and Gng8 across both the apical and basal populations of neuronal precursors.

